# The transcription factor Osr1 regulates epithelial-mesenchymal crosstalk during embryonic bladder development

**DOI:** 10.1101/2025.10.10.680757

**Authors:** V Murugapoopathy, IR Gupta

## Abstract

The molecular events that define cell fate decisions during bladder development are poorly characterized. Here, we establish a temporal single-cell atlas when the bladder first arises from the cloaca until its major layers have been established that include the uroepithelium, the lamina propria and the smooth muscle. The analysis resolved the cell origin of ligands and their respective receptors for four major signaling pathways that have been previously implicated in bladder development, SHH, BMP, WNT and FGF. The transcription factor Odd-skipped related 1 is essential for mesenchymal differentiation during organogenesis of the foregut, kidney, limb, ureter and is highly expressed during bladder development. We demonstrate that Osr1 is required for development of the bladder: homozygous loss of *Osr1* results in depletion of smooth muscle, loss of extracellular matrix, loss of suburothelial cells, and a less stratified epithelium lacking intermediate and superficial cells. Transcripts within the four major signaling pathways, SHH, BMP, WNT and FGF, were decreased during cellular diversification in bladders from *Osr1* homozygous null embryos. In summary, Osr1 is a central mediator of epithelial-mesenchymal crosstalk and cell fate decisions during bladder development.

## Introduction

The bladder forms as a distinct outgrowth from the cloaca to allow for storage and expulsion of urine at appropriate times. Bladder dysfunction affects 20 % of children after the age of 5 years (Fuentes et al., 2019) and up to 50% of adults over the age of 65 (Reynolds et al., 2016) and is associated with recurrent urinary tract infections, upper urinary tract dilatation, and chronic kidney disease. Despite the significant psychosocial, economic, and medical burden of bladder dysfunction, effective therapies outside of catheterization do not exist and augmentation of the bladder often uses gut-derived tissue which is more permeable resulting in a higher risk of bladder infections, metabolic derangements, and vitamin deficiencies. Bladder organoid models have been primarily used in cancer research but have been limited by a lack of knowledge of the cell types and signaling events required for bladder formation.

The bladder stores and expels urine by coordination and communication between the epithelium, lamina propria and smooth muscle layers. The epithelium consists of 3 cell layers: the basal and intermediate layers which harbor progenitor cells that can restore the epithelium (Wiessner et al., 2022) and the superficial layer which serves as a barrier to urine and pathogens in the bladder lumen (Lavelle et al., 2002). The lamina propria contains large amounts of extracellular matrix proteins like collagen I, collagen III, elastin and fibronectin that bear most of the mechanical load during distension (Andersson & McCloskey, 2014b). Cells in the lamina propria, primarily within the suburothelial region, relay messages between the epithelium and muscle during filling and emptying of the bladder (Fry et al., 2007). The smooth muscle layer consists of longitudinal and circular bundles that contract the bladder from all directions to to allow complete emptying (Chang et al., 1999).

During development, the epithelium forms from a single layer as an outgrowth from the cloaca and invades the surrounding tail bud mesenchyme which gives rise to the future lamina propria and smooth muscle (Georgas et al., 2015; Tasian et al., 2010). Crosstalk between the epithelium and lamina propria is essential for patterning both layers. Sonic hedgehog (SHH) is secreted from the epithelium and forms a gradient from high to low across the mesenchyme. Low levels of SHH permit smooth muscle formation in the outer mesenchyme (M. Cao et al., 2010; DeSouza et al., 2013; Haraguchi et al., 2007), while high levels permit the lamina propria to form adjacent to the epithelium. Other signaling pathways involved in bladder development include the FGF, WNT and BMP signaling pathways. FGFR2 patterns the mesenchyme (Ikeda et al., 2017) and Wnt3 is required for cloacal development in zebrafish (Baranowska Körberg et al., 2015). Bmp4 regulates terminal differentiation of the uroepithelium and may be implicated in smooth muscle differentiation, however its role is poorly understood (Islam et al., 2013; Mysorekar et al., 2009; C. Wang et al., 2017).

Transcription factors coordinate inputs from multiple signaling pathways including SHH, BMP, FGF and WNT, to regulate cell fate decisions. The zinc finger transcription factor, Odd-skipped related 1, Osr1, functions downstream of the SHH pathway and also binds to the Bmp4 promoter to repress Bmp4 expression. The SHH-BMP4 signaling axis and Osr1 have been implicated in mesenchymal cell fate decisions in the lungs, ureter, and foregut (Bohnenpoll et al., 2017; Bottasso-Arias et al., 2022; Han et al., 2017; Rankin et al., 2012; Straube et al., 2025), suggesting that Osr1 may coordinate the SHH-BMP4 signaling axis (Brenner-Anantharam et al., 2007; M. Cao et al., 2010; Han et al., 2017; Rankin et al., 2012). Previously, we found that Osr1 is strongly expressed in the precursor to the bladder, the urogenital sinus, at embryonic day 11 in the mouse and throughout bladder development (Fillion et al., 2017). Heterozygous loss of *Osr1* resulted in defects in the bladder mesenchyme including loss of collagen I and a decrease in suburothelial cells at the newborn stage (Murugapoopathy et al., 2021) that resulted in decreased bladder capacity in the adult. These observations led us to hypothesize that Osr1 regulates mesenchyme differentiation in the bladder by coordinating the SHH-BMP4 signaling axis.

## Results

### Osr1 is expressed in mesenchymal progenitor cells and their derivatives

To understand the timing and cell fate trajectories during bladder mesenchymal differentiation, we performed the first single-cell transcriptomic analysis of the embryonic mouse bladder. Three timepoints were examined: embryonic day (E) 12, when the bladder becomes a distinct outgrowth from the cloaca, E14, when smooth muscle differentiation begins, and E15, when the mesenchyme forms distinct compartments of inner and outer lamina propria and smooth muscle (Georgas et al., 2015). At E12, adder is not externally visible under the microscope, therefore, the tissue above the genital tubercle and beneath the umbilical artery was dissected and separated from the hindgut tube (Figure 1A). At E14 and E15, the bladder is directly underneath the skin above the genital tubercle (Figure 1A). Bladders were dissected and pooled from 2 embryos/timepoint (Figure 1A). After filtering out dying cells (high mitochondria RNA content), cells with low transcript number, and doublets, transcriptomes from 6069, 8392, and 8471 cells for E12, E14, and E15 bladders, respectively, were integrated using Seurat and Harmony packages in R (Hao et al., 2024; Korsunsky et al., 2019) (Figure 1C). After unsupervised clustering, 9 cell types were identified with the majority representing mesenchymal progenitors and their derivatives (Figure 1B, D). Cell types were characterized using canonical markers (Figure 1E) (Yu et al., 2019). To determine the developmental trajectories of mesenchymal progenitors, mesenchymal cells were sub-clustered (Supplemental Figure 1), and analyzed using Monocle3 (Figure 1F) (J. Cao et al., 2019). At E12, the mesenchyme was largely undifferentiated with 2 clusters of proliferating and non-proliferating mesenchymal progenitors observed that expressed *Top2a* and *Pdgfra+*, respectively (Figure 1E,G). At E14, smooth muscle and fibroblast cells were detected and likely arose from a common progenitor as seen by pseudotime lineage analysis (Figure 1 D,F). Finally, at E15, most of the mesenchymal progenitors differentiated to smooth muscle and fibroblast cell fates based on their increased expression of *Acta2* and *Col3a1*, respectively (Figure 1D,E). Confirming our previous characterization (Fillion et al., 2017; Murugapoopathy et al., 2021), the transcription factor Osr1 was highly expressed in mesenchymal progenitors as well as their derivatives (Figure 1H). Spatially, we observed Osr1mRNA expression in all layers and at all timepoints suggesting that Osr1 may regulate cell fate decisions during bladder development (Figure 1I).

**Figure 1.**
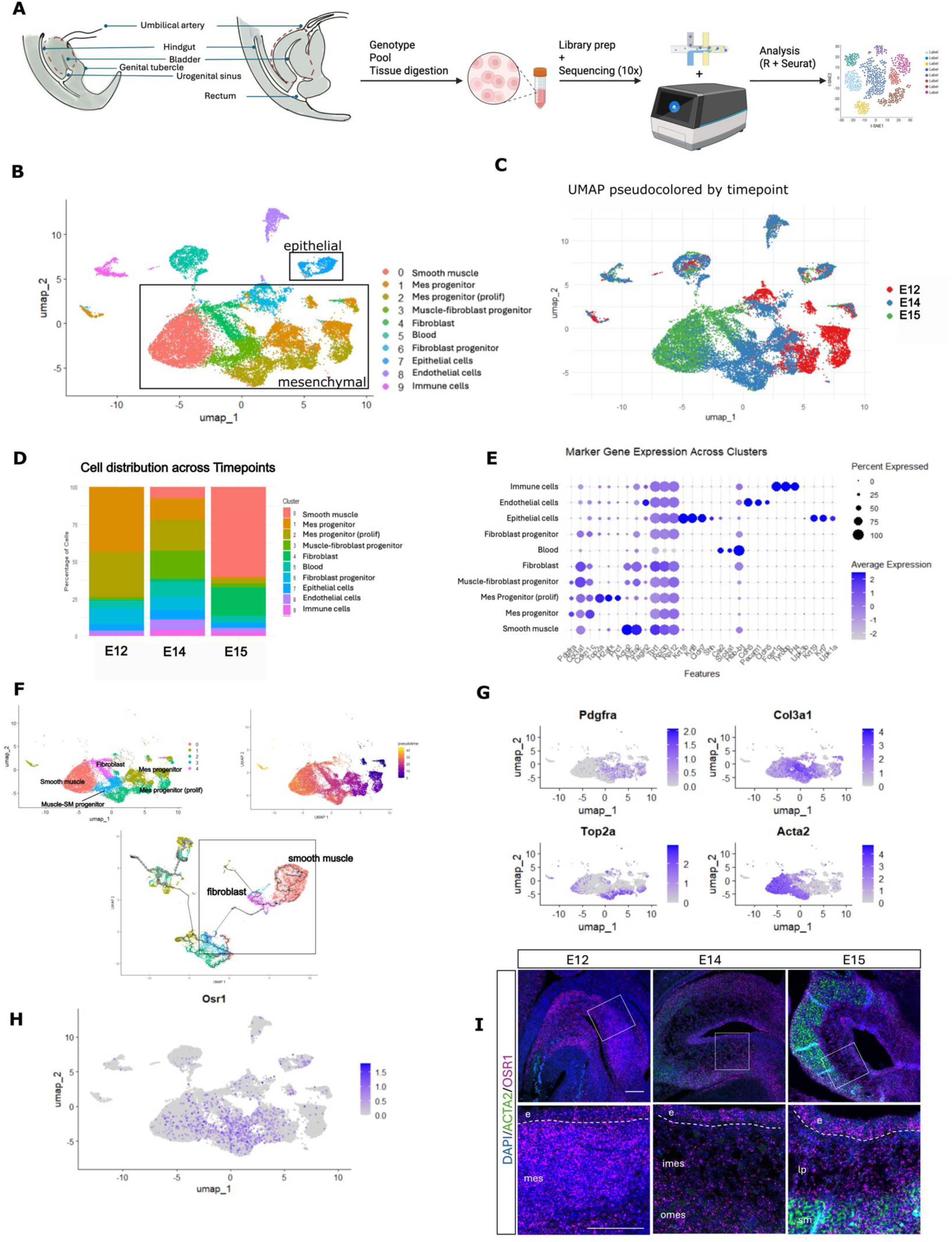
Osr1 is expressed in mesenchymal progenitor cells and their derivatives. (A) Embryonic wildtype (WT) bladders were dissected from the regions depicted (dotted lines) at an early timepoint (embryonic day 12 (E12), and later timepoints (E14 and E15 shown as one cartoon), dissociated into single cell suspensions followed by 10X library preparation and sequencing. (B) UMAP visualization and unsupervised clustering of E12, E14, and E15 integrated data. (C) Pseudocoloring of timepoints demonstrates the transition of cells from primarily progenitor-like states at E12 towards more differentiated cell types at E14 and E15. (D) Distribution of cell types across timepoints. (E) Clusters identified using canonical markers. (F) Pseudotime trajectory analysis of mesenchymal ell populations reveals a common intermediate precursor (light blue) that differentiates into either smooth muscle (red) and fibroblast cells (pink) (boxed). (G) Visualization of markers for different mesenchymal cell types. Mesenchymal progenitors and their derivatives including fibroblasts showed high expression of *Pdgfra*. Differentiated fibroblasts expressed high levels of *Col3a1*. Muscle cells expressed *Acta2* but no *Pdgfra*. *Top2a* was seen in mesenchymal progenitor cells undergoing proliferation at E12 and E14. (H,I) *Osr1* expression was expressed in mesenchymal and epithelial cell types across all 3 timepoints and validated by fluorescent *in situ* RNA hybridization (*Osr1* signal in pink and *Acta2* signal in green). mes – mesenchyme, imes – inner mesenchyme, omes – outer mesenchyme, e – epithelium, sm – smooth muscle, lp – lamina propria. Scale bar = 100 um.

### Resolution of BMP, FGF, HH and WNT signaling pathways that regulate bladder mesenchyme differentiation

BMP, FGF, HH, and WNT signaling are required during bladder development, however the specific ligands, receptors, and cell types by which they exert their effects remain largely unresolved (Baranowska Körberg et al., 2015; M. Cao et al., 2010; Ikeda et al., 2017; Islam et al., 2013). The single-cell RNA sequencing analysis revealed that *Bmp4* is the primary BMP ligand expressed in the bladder, primarily from mesenchymal progenitor cells at E12 and E14 and fibroblasts (Figure 2A) but not differentiated smooth muscle cells. BMP4 signals to its receptors *Bmp1ra* and *Bmpr2* that are expressed in both mesenchymal and epithelial cells suggesting that BMP signaling may be important for mesenchyme and epithelial patterning and crosstalk between the layers during bladder development (Figure 2E). Mesenchymal progenitors also expressed high levels of *Fgfr1* (Figure 2B,F) which may respond to the ligand *Fgf9* expressed by the mesenchyme (Figure 2B). The epithelium, in contrast, expressed high levels of *Fgfr2*. Interestingly, a role for Fgfr2 has been characterized in the bladder mesenchyme but not the bladder epithelium (Ikeda et al., 2017) SHH is secreted from the bladder epithelium (Figure 2C) and mesenchymal progenitors expressed high levels of *Ptch1* (Figure 2C,G). Unlike Bmp1ra and Fgfr1, *Ptch1* expression was decreased in differentiated mesenchymal progenitors that became fibroblast and smooth muscle (Figure 2G). In terms of the WNT signaling pathway, mesenchymal progenitors at E12 primarily produced *Wnt5a*, while the muscle-fibroblast progenitors and fibroblasts expressed *Wnt2* (Figure 2D). Epithelial cells primarily expressed *Wnt4*. The mesenchymal progenitor cells expressed high levels of *Fzd1* and *Fzd2*, with differentiated smooth muscle and fibroblasts expressing lower levels of both receptors when compared to progenitor cells (Figure 2H). This suggests that WNT signaling may pattern the mesenchyme and regulate mesenchymal-epithelial crosstalk. The undifferentiated mesenchyme, present primarily at E12 (Clusters 1&2), uniquely produced *Wnt5a* and *Fgf9* (Figure 2B,D), which were no longer expressed in the more differentiated cell types suggesting these ligands may be important during the pre-differentiation stages of bladder development. The cell origin of ligands is summarized in Figure 2I, with the mesenchymal cells producing Bmp4, Fgf9, Wnt5a and Wnt2 and the epithelia producing Shh and Wnt4

**Figure 2.**
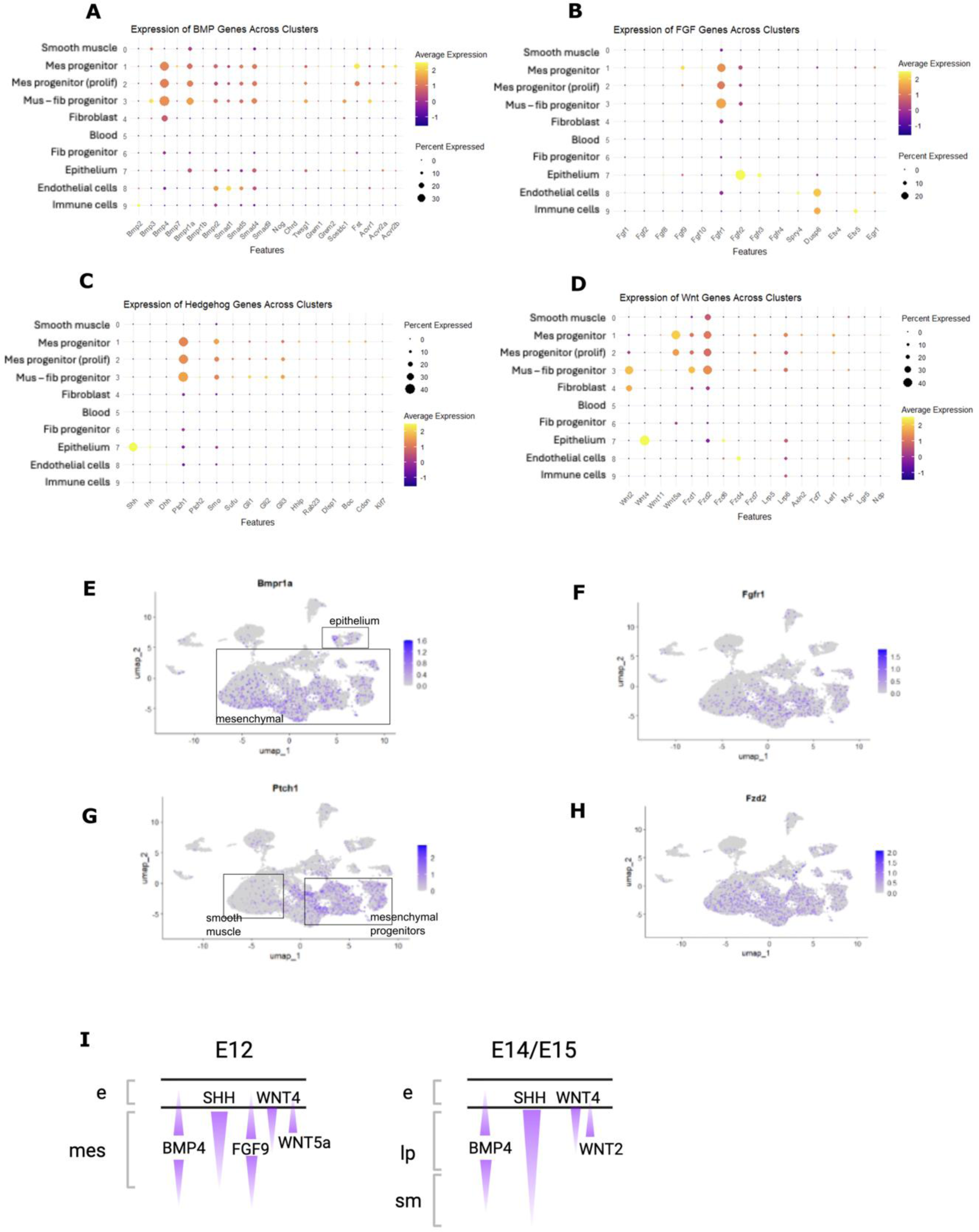
Resolution of BMP, FGF, HH and WNT signaling pathways that regulate bladder mesenchyme differentiation. (A,E) Expression of BMP genes across clusters revealed high levels of *Bmp4* in mesenchymal progenitors and their derivatives except in smooth muscle cells. Fibroblasts and mesenchyme progenitors, but not smooth muscle cells, also expressed high levels of the *BMP* receptor *Bmp1ra* along with the downstream transcription factors *Smad1*, *5* and *4*. (B,F) FGF pathway analysis revealed high levels of *Fgfr1* expressed on mesenchymal cells while epithelial cells expressed high levels of *Fgfr2*. The main detectable FGF ligand was *Fgf9,* secreted from mesenchymal progenitors. (C,G) HH pathways analysis confirmed release of SHH from the epithelia and high levels of its receptor *Ptch1* expressed in mesenchymal progenitors with low expression in differentiated smooth muscle. This corresponded with expression of the transcription factor *Gli3*. (D,H) Analysis of Wnts revealed high levels of *Wnt2* secreted by fibroblasts and muscle-fibroblast progenitors, while uncommitted mesenchymal progenitors expressed high levels of *Wnt5a*. In contrast, epithelial cells secreted *Wnt4*. Mesenchymal progenitors expressed the receptors *Fzd1* and *Fzd2*, while the epithelium expressed *Fzd6*. (I) Summary of cell origin of ligands and directionality of their signaling pathways. e- epithelium, mes- undifferentiated mesenchyme, lp- lamina propria, sm- smooth muscle.

To determine if Osr1 interacted with any of these signaling pathways, two published ChIP sequencing datasets in mouse nephron progenitor cells and in chick limb mesenchyme were re-analyzed to look for common and tissue-specific targets of Osr1 (Table 1) (O’Brien *et al*., 2018; Orgeur *et al*., 2018). Osr1 bound to promoters for the receptors *Bmpr1b* and *Fgfr2* in both datasets. Osr1 also bound to *Fgfr1*, *Ptch2* and *Fzd2* in the mouse and *Ptch1* and *Fzd1* in the chick. Osr1 also bound to *Wnt5a* and *Fgf9* in the mouse kidney, which are both expressed in the early E12 mesenchymal progenitors in the mouse bladder. Taken together, Osr1 likely binds to the promoter regions of ligands and receptors within the BMP, FGF, WNT, and SHH pathways in the bladder epithelia and mesenchyme during bladder development.

**Table 1:** Cross-analysis of CHIP-seq datasets (O’Brien et al., 2018; Orgeur et al., 2018) CHIP seq analysis revealed several binding sites for Osr1 in the pathways analyzed with overlap between targets in the kidney and limb suggesting likely interaction of Osr1 with these targets in the bladder as well. Smad4, Fgfr2, Fgfr1, Fgf9, Gli2, Gli3, Ptch1, Smo, Wnt4, Wnt5a, Fzd1, and Fzd2 are all expressed in the mouse embryonic bladder.

| Pathway | Genes in common | Genes unique to mouse nephron progenitors | Genes unique to chick limb mesenchyme |
| --- | --- | --- | --- |
| BMP | Tgfr3, Inhba, Ltbp2, Bmpr1b | Bmp7, Bmp6, Bmp2, Smad2 | Bmp5, Bmp8, Smad4 |
| FGF | Fgfr2 | Fgfr1, Fgf9 | Fgf4, Fgf16 |
| Hedgehog | Gas1, Gli2 | Ptch2, Smo | Gli3, Ptch1 |
| Wnt | Wnt4, Wnt11, Dact2, Bcl9l | Wnt10a, Wnt5a, Wnt7b, Fzd10, Fzd2 | Fzd1 |

### Loss of Osr1 impairs initiation of smooth muscle differentiation

We previously reported that heterozygous loss of *Osr1* caused a reduction in extracellular matrix and suburothelial cells in the lamina propria, as well as bladder dysfunction at the adult stage (Murugapoopathy et al., 2021). The defect in bladder mesenchyme from haploinsufficiency of *Osr1* coupled with the strong expression of *Osr1* in mesenchymal progenitors during bladder development prompted us to examine cell differentiation with and without *Osr1*. We performed single-cell RNA sequencing on *Osr1* KO and WT embryonic bladders at E12, E14 and E15. Cell numbers and cell viability prior to sequencing are summarized in Supplemental Table 1. *Osr1*-KO embryos do not survive after E15 and only 25% of KO embryos survive between E12-15 (Supplemental Figure 3). At E12, *Osr1-*KO cells, pooled from 2 bladders (n=2672 cells after quality control and pre-processing), were integrated with WT cells, also pooled from 2 bladders (n=4103 cells after quality control and pre-processing). We noted decreased cell viability in E12 KO bladder cells that may explain the lower cell numbers. However, TUNEL analysis did not reveal increased cell death in the putative bladder (Supplemental Figure 3). As shown (Figure1A), it is likely that non-bladder tissue was included during cell collection. After integration and neighborhood assignment, thirteen clusters were identified through unsupervised clustering. There were no large differences in the cell types present between WT versus KO bladders and most mesenchymal cells were undifferentiated. The KO bladders showed a larger proportion of fibroblast/fibroblast progenitor cells and decreased early mesenchymal progenitors and muscle progenitors suggesting a shift towards fibroblast cell fates (Figure 3A). At this stage, bladder morphology and tissue architecture were similar in WT and KO bladders: the bladder mesenchyme appeared homogeneous without a distinct muscle or lamina propria layer (Figure 3B). To determine if there were transcriptional differences between WT and KO bladders prior to cell differentiation, we performed pseudo-bulk analysis followed by GO term enrichment (Figure 3C, D). Interestingly, KO embryos had a significant decrease in H2A.Z transcripts (Figure 3C), a histone variant that modulates chromatin accessibility and that has previously been shown to interact with Osr1 (Zhang et al., 2017). There were also decreased levels of genes associated with RNA splicing and cell proliferation suggesting that bladder is affected. There was increased expression of genes associated with extracellular matrix like *Col1a1* supporting the shift in cell fate towards fibroblasts in KO bladders. To determine if there were differences in the signaling pathways previously analyzed (Figure 2), we compared WT and KO transcript levels after pseudo-bulk analysis. This revealed that there was significantly lower expression of, *Smo*, and *Wnt5a* in the KO bladders, with an overall trend showing a decrease in components of the BMP and HH pathways. This could be due to a lack of Osr1 binding to the regulatory sites of these genes as observed in the re-analyzed ChIP-sequencing datasets (Table 1). The only significantly upregulated gene in KO bladders at E12 was *Wnt2*.

**Figure 3.**
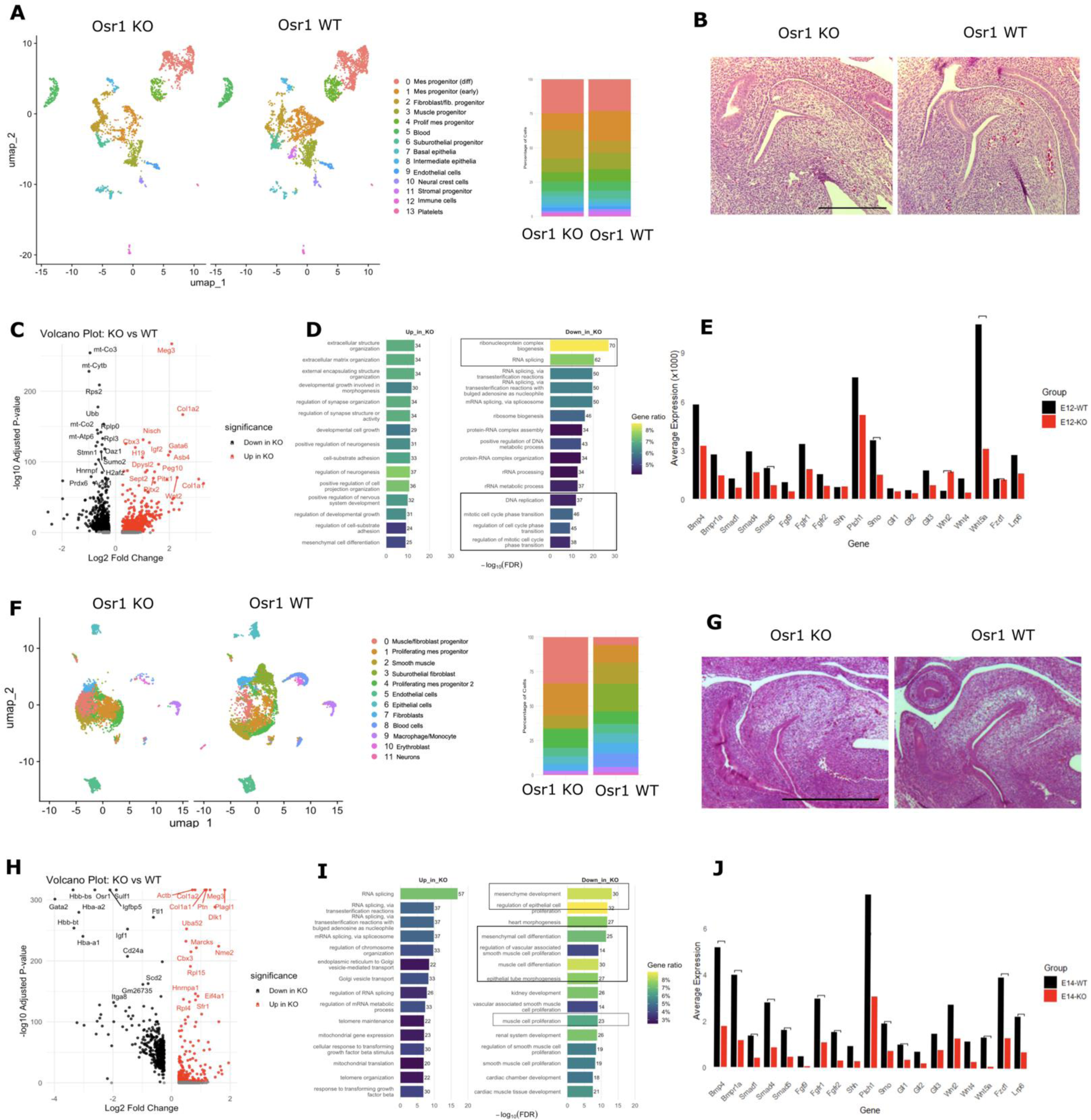
Loss of Osr1 impairs initiation of smooth muscle differentiation. (A) Comparative UMAPs of WT and KO embryonic bladder cells at E12 revealed differences in distribution of mesenchymal progenitor cells with less early progenitors (cluster 1) and more fibroblast progenitors (cluster 2) seen in KO bladders. (B) No major differences in histology between WT and KO bladders were seen. (C,D) Differential gene expression revealed upregulation of *Col1a1* and *Col1a2* in KO bladders while GO term analysis suggested major differences in cell cycle and proliferation pathways (boxed). (E) Analysis of signaling pathways identified significant downregulation of *Smad5*, *Smo*, and *Wnt5a* in KO bladders. Brackets indicate adjusted p value < 0.05 (F, G) At E14, differences were seen in cell differentiation of KO bladders with failure to form smooth muscle and cells trapped in mesenchymal progenitor-like states which was confirmed by H&E staining.(H,I) Differential expression analysis revealed differences in mesenchymal differentiation pathways and smooth muscle development (boxed). (J) Major differences were seen in signaling pathways including significant downregulation of *Bmp4* and its receptor *Bmp1ra*, *Smad1*, *Smad5*, *Smad4*, *Fgfr1* and *Fgfr2*, *Smo* and *Gli1*, and *Wnt5a, Fzd1, and Lrp6* in KO bladders. Brackets indicate adjusted p value < 0.05.

The single-cell sequencing data at E14 (8850 WT cells, and 6616 KO cells) revealed marked changes in the abundance of cell clusters with a loss of smooth muscle, fibroblast progenitor cells and suburothelial fibroblasts in KO bladders (Figure 3F). The majority of mesenchymal cells were trapped in a muscle-fibroblast progenitor-like state, suggesting differentiation was impaired in KO bladders. This was also supported by histology that showed a diminished smooth muscle layer in KO bladders as seen by H&E staining (Figure 3G). At E14, the KO bladders showed significant downregulation of genes and pathways involved in mesenchymal cell differentiation including *Gata2* which has previously been shown to be important for urogenital mesenchyme differentiation (Khandekar et al., 2004) as indicated by the GO term analysis (Figure 3 H,I). There continued to be increased expression of extracellular matrix proteins such as *Col1a1* in the KO bladders. In contrast to E12, genes associated with RNA splicing and cell proliferation were now upregulated, which could suggest that these processes were delayed, but not completely perturbed with loss of *Osr1*. We observed similar levels of proliferation in WT and KO bladders using Ki67 immunofluorescent staining (Supplemental Figure 4). Consistent with the GO-term analysis indicating major loss of mesenchymal and epithelial differentiation pathways, targeted inspection revealed that *Bmp4*, *Bmp1ra*, *Smad1*, *Smad 5*, Smad 4, *Fgfr1*, *Fgfr2*, *Smo*, *Wnt5a* and *Fzd1* were all significantly downregulated in KO bladders (Figure 3J).

### Loss of Osr1 results in reduced smooth muscle, suburothelial cells, and collagen I in the differentiated bladder

We integrated 8471 WT cells and 6237 KO cells for analysis at E15, the latest timepoint at which *Osr1*-KO embryos are viable. By E15, most mesenchymal progenitors differentiated to become smooth muscle or fibroblast cells in WT bladders, as shown in both the single-cell sequencing data and histology (Figure 4A, B). In the KO embryos, the bladder was more oblong in shape, and the wall thickness was significantly reduced due to the thin underdeveloped smooth muscle. In contrast, the WT bladders had a thick smooth muscle layer with circular and longitudinal muscle bundles (Figure 4B). There was also a distinct difference in lamina propria development: the WT bladders had a cell-dense suburothelial layer of cells, while the KO bladders had a more uniform lamina propria without a discrete suburothelial layer (Figure 4B). From the single-cell sequencing data, there was an increased number of cells arrested in a mesenchymal progenitor-like state in the KO bladders similar to what was seen at E14 (Figure 4A). From the pseudobulk DEG and GO analysis, an upregulation of *Gata2* was observed in KO bladders, in contrast to E14, suggesting a potential recovery of mesenchymal cell differentiation (Figure 4C, D). GO analysis revealed decreases in cellular respiration and metabolic processes that are typically associated with proliferation and differentiation as well as downregulation of muscle cell differentiation suggesting an incomplete recovery of muscle differentiation pathways (Figure 4D). Meanwhile pathways involved in stem cell maintenance were upregulated. Targeted analysis of signaling pathways was noteworthy in that the pathways that had been downregulated at E14 including *Bmp4*, Fgfr1, Smo, and *Fzd1*, were now all significantly increased (Figure 4E). However, *Bmp1ra* remained significantly decreased. Finally, *Shh* and its targets *Gli1*, *Gli2*, and *Gli3* were expressed at similar levels in WT and KO. This recovery and increase in components of the Bmp and Wnt signaling pathways suggested a compensatory response. Because Osr2 can compensate for loss of Osr1, transcript levels of *Osr2* were examined at the three timepoints (Gao et al., 2009; Neto et al., 2012; Rankin et al., 2012). Indeed, there was a significant increase in *Osr2* transcript levels at E15 (Figure 4F). To define the location of the increased number of mesenchymal progenitors, bladder sections were assessed for vimentin expression, which detects mesenchymal progenitors and fibroblasts, but not smooth muscle (Council & Hameed, 2009). An increased number of vimentin-positive cells was seen in the smooth muscle layer of *Osr1*-KO when compared to WT bladders by immunofluorescent staining (Figure 4G). To confirm that these cells were mesenchymal progenitors and not differentiated fibroblasts, collagen I expression was examined. In contrast to the WT bladders that showed robust collagen I deposition in the lamina propria with weak expression in the muscle layer, there was almost no collagen expression in the KO bladders with only weak expression within the lamina propria layer.

**Figure 4.**
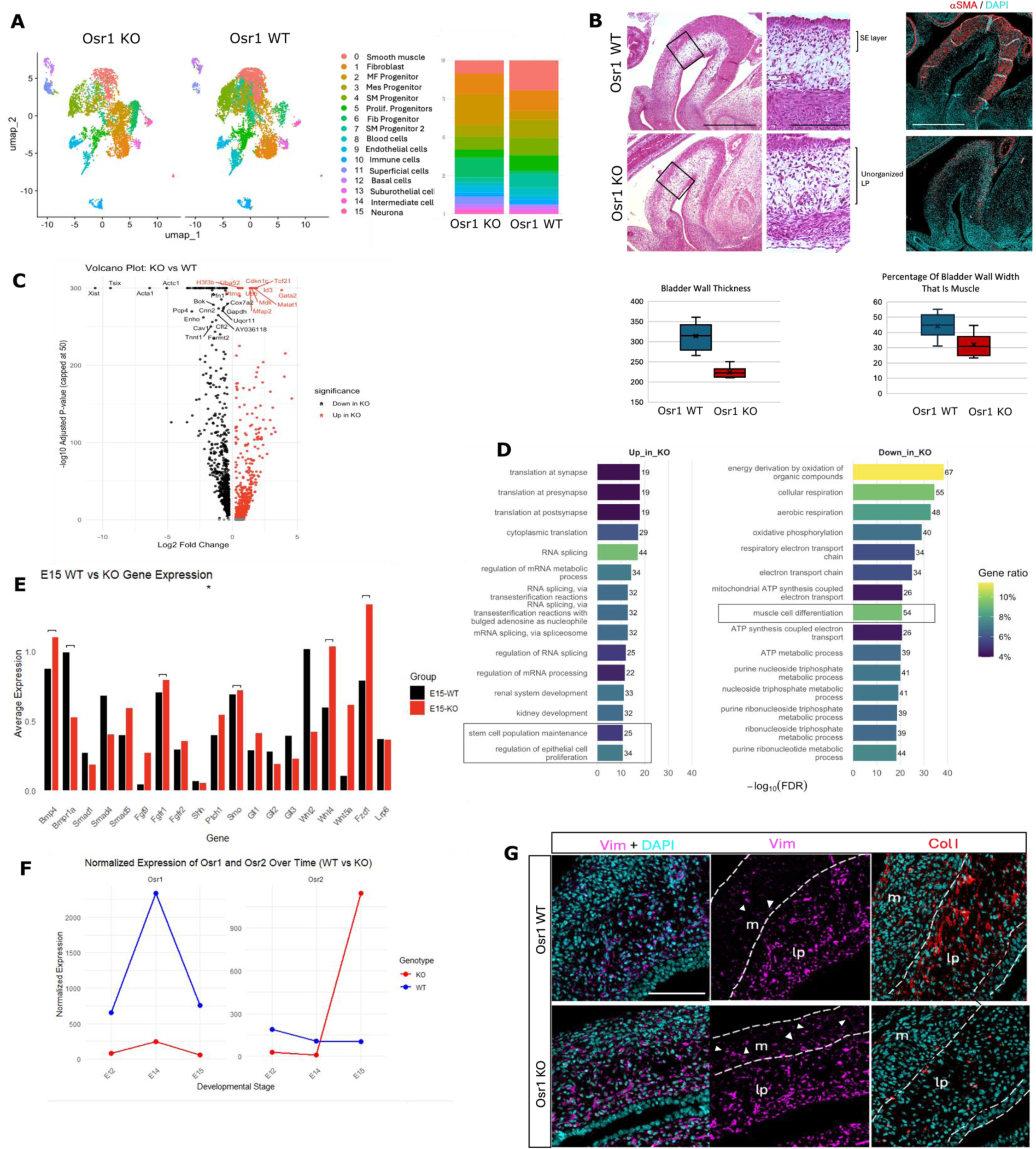
Loss of Osr1 results in reduced smooth muscle, suburothelial cells, and collagen I in the differentiated bladder. (A) Comparative UMAPs of WT and KO embryonic bladder cells at E15 revealed decreased smooth muscle cells and continued arrest of cells in mesenchymal progenitor like states. (B) *Osr1-* KO bladders had reduced thickness (bladder wall thickness) from a depleted smooth muscle layer (percentage of bladder wall) that corresponded with decreased aSMA protein expression compared to WT bladders. The lamina propria of the KO bladders also exhibited a loss of a discrete suburothelial population which was seen in the WT bladder. Scale bar = 500 um for whole bladder and 100 um for inset. (C,D) Differential gene expression showed significant differences in muscle and mesenchymal differentiation seen in GO term analysis (boxed) with downregulation of *Acta1 in KO* bladders seen in the volcano plot. Meanwhile pathways involved with stem cell maintenance were upregulated (boxed). (E) Gene expression comparison showed increased expression of *Bmp4*, *Fgfr1*, *Smo*, *Wnt4,* and Fzd1 in KO bladders and reduced expression of *Bmp1ra*. (F) *Osr2* expression was increased at E15 in the KO possibly as a compensation for the loss of *Osr1* (G) Immunofluorescent detection showed increased vimentin expression (magenta) in the smooth muscle and significant depletion of collagen I (red) in KO bladders, suggesting fibroblasts are unable to secrete collagen. m – muscle, lp – lamina propria, arrows – vimentin positive cells in muscle layer. Scale bar = 100 um.

Although there were defined fibroblast populations and upregulation of collagen transcripts in the KO mice (Cluster 1, Figure 4A, 3H) the fibroblasts were unable to secrete collagen. Taken together, our data shows that loss of *Osr1* results in severe defects in bladder mesenchymal development with a smaller bladder wall, a reduction in the smooth muscle layer that contains more mesenchymal progenitors, and a reduction in suburothelial cells of the lamina propria.

### Loss of Osr1 results in defects in bladder epithelial stratification

Throughout bladder development, *Osr1* transcripts were detected in the epithelium (Figure 1H, I). Epithelial stratification begins at E14 in the bladder and by E15, three layers have formed: basal, intermediate and superficial. Inspection of *Osr1-*KO bladders at E15 revealed that the epithelium was less stratified (Figure 4G, Figure 5C). Because of the smaller number of epithelial cells, the single cell sequencing data from WT and KO bladders was pooled by combining E14 and E15 timepoints and yielded 513 WT and 269 KO epithelial cells. The WT and KO datasets were extracted and sub-clustered using canonical markers (Jafari & Rohn, 2022; Wiessner et al., 2022)(Figure 5 A,B). The single cell sequencing data combined with immunofluorescent staining for keratin proteins revealed a failure of epithelial stratification in KO bladders (Figure 5 A, E). Most epithelial cells in the KO bladder were basal cells with fewer intermediate and superficial cells compared to the WT bladder (Figure 5A). Differential gene expression revealed a decrease in genes associated with umbrella cells in the superficial layer including *Upk3a* and *Upk3b.* This correlated with the GO terms that showed downregulation of processes like urogenital system development, suggesting an overall decrease in epithelial differentiation (Figure 5 C, D). Epithelial cells expressed receptors for Wnt (*Fzd2/Fzd6*), FGF (*Fgfr2*), and BMP (*Bmp1ra*) signaling (Figure 2). All of these pathways were depleted at E14 in KO bladders demonstrating that Osr1 regulates epithelial differentiation and growth (Figure 4J).

**Figure 5.**
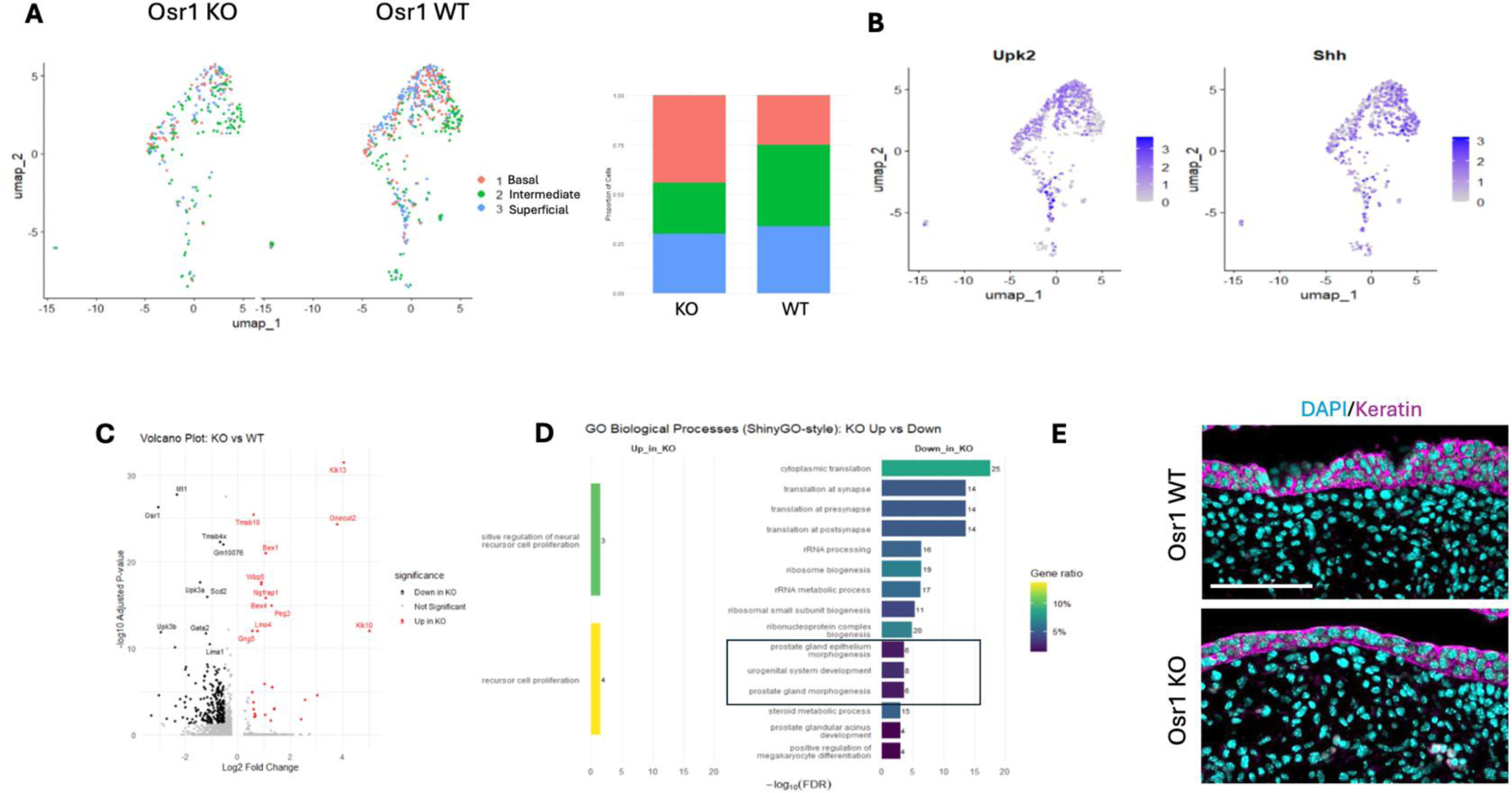
Loss of Osr1 results in defects in bladder epithelial stratification. WT and KO epithelial cells from E14 and E15 were isolated from the dataset through expression of canonical markers and subclustered based on expression of previously characterized markers including Shh and Upk2: basal (Shh-high, Upk-negative), superficial cell (Upk2-high, Shh-negative), and intermediate cells (Shh-positive, Upk-positive) (A, B) Distribution of cell types was skewed towards basal cells in the KO. (C) Differential gene expression analysis with upregulated genes (red) and downregulated genes (black) in the KO bladders revealed a decrease in umbrella cell genes such as *Upk3a*. (D) GO Biological Process analysis revealed disruption of epithelial morphogenesis pathways (boxed). (E) Depletion of the epithelial layer was demonstrated by labeling keratin proteins in E15 bladders. Scale bar = 100 um.

### Hedgehog and BMP signaling are essential for patterning the bladder mesenchyme

SHH and BMP signaling have previously been shown to be part of a gene regulatory network that coordinates mesenchymal cell fate determination, smooth muscle differentiation and mesenchyme patterning in the gut, ureter, and bladder (Bohnenpoll et al., 2017; Cheng Wei et al., 2008; Huycke et al., 2019). The single cell RNA sequencing data demonstrated a decrease in *BMP4* and its receptor *BMP1ra*, as well as a decrease in the HH genes, *Smo* and *Gli1*, that corresponded to a depletion of the smooth muscle and lamina propria in E14 KO bladders (Figure 3J). To distinguish the effects of these pathways, bladder explants were cultured in media containing inhibitors for Alk2/3 (also known as BMP1ra) (LDN193189) and Smo (Sonidegib/LDE225), which inhibit the BMP and HH pathways, respectively, or a vehicle (DMSO) control at E13.5. Concentrations of inhibitors were selected based on previous literature (Cross et al., 2011; Graziosi et al., 2024; Hadas et al., 2024; Skvara et al., 2011). Bladder explants were cultured for 72 hours and growth was evaluated (Supplemental Figure 5) and sections were processed for histology and immunofluorescence. Bladders treated with BMP inhibitor were similar in size to controls, while those treated with HH inhibitor trended towards being smaller (p= 0.09) (Figure 6A-C, G). We then analyzed the percentage of the bladder wall that was muscle and used expression of alpha smooth muscle actin to evaluate muscle differentiation. BMP-inhibited bladders had significantly decreased smooth muscle thickness (p = 0.02) and a trend towards decreased alpha smooth muscle actin expression (Figure 6 A’, B’, H, I). HH-inhibited bladders showed a reduced lamina propria layer and trended towards an increase in smooth muscle and expression of alpha smooth muscle actin (Figure 6 A’, C’, H, I). We next evaluated whether mesenchymal progenitors were trapped in progenitor-like states or shifted towards fibroblast-like states by staining for vimentin. BMP-inhibited bladders exhibited increased vimentin expression in the lamina propria when compared to controls, potentially indicating an expansion of the fibroblast population (Figure 6 A’’, B’’, I) (p= 0.03). Similar to the *Osr1*-KO group at E15, BMP-inhibited bladders also had increased vimentin-positive cells in the smooth muscle compartment (p = 0.0004), suggesting that mesenchymal cells in the smooth muscle layer were either trapped in progenitor-like states or shifted towards fibroblast-like states. SHH-inhibited bladders also showed increased vimentin expression in both the lamina propria and smooth muscle layers (p = 0.04, 0.02) (Figure 6 A’’, C’’, I). Finally, to determine if BMP and/or SHH inhibition was able to replicate the loss of collagen seen in the *Osr1*-KO bladders, we examined COL1A1 expression in the lamina propria and smooth muscle layers. Surprisingly, the BMP inhibition group showed strikingly elevated levels of collagen deposition throughout the lamina propria and smooth muscle layers (p = 0.01, 0.01) suggesting a shift in mesenchymal cell fate towards fibroblasts (and away from smooth muscle), an increased production of extracellular matrix, and/or a combination of both (Figure 6 D, E, J). In contrast, HH-inhibited bladders did not show significant changes in collagen expression in either the lamina propria or smooth muscle, suggesting that the increased vimentin-positive cells were likely trapped as progenitors, as opposed to shifted towards fibroblasts. Neither SHH nor BMP inhibition was able to replicate the collagen loss seen in the *Osr1*-KO bladder suggesting that loss of collagen is via another pathway. Taken together, SHH inhibition was able to replicate the smaller bladders seen in the *Osr1*-KO bladders, while BMP inhibition replicated the depletion of the smooth muscle layer. Both SHH and BMP inhibition resulted in increased vimentin-positive cells in the lamina propria, but the increased collagen seen with BMP inhibition suggests that loss of BMP signaling shifts mesenchymal progenitors towards fibroblast fates.

**Figure 6:**
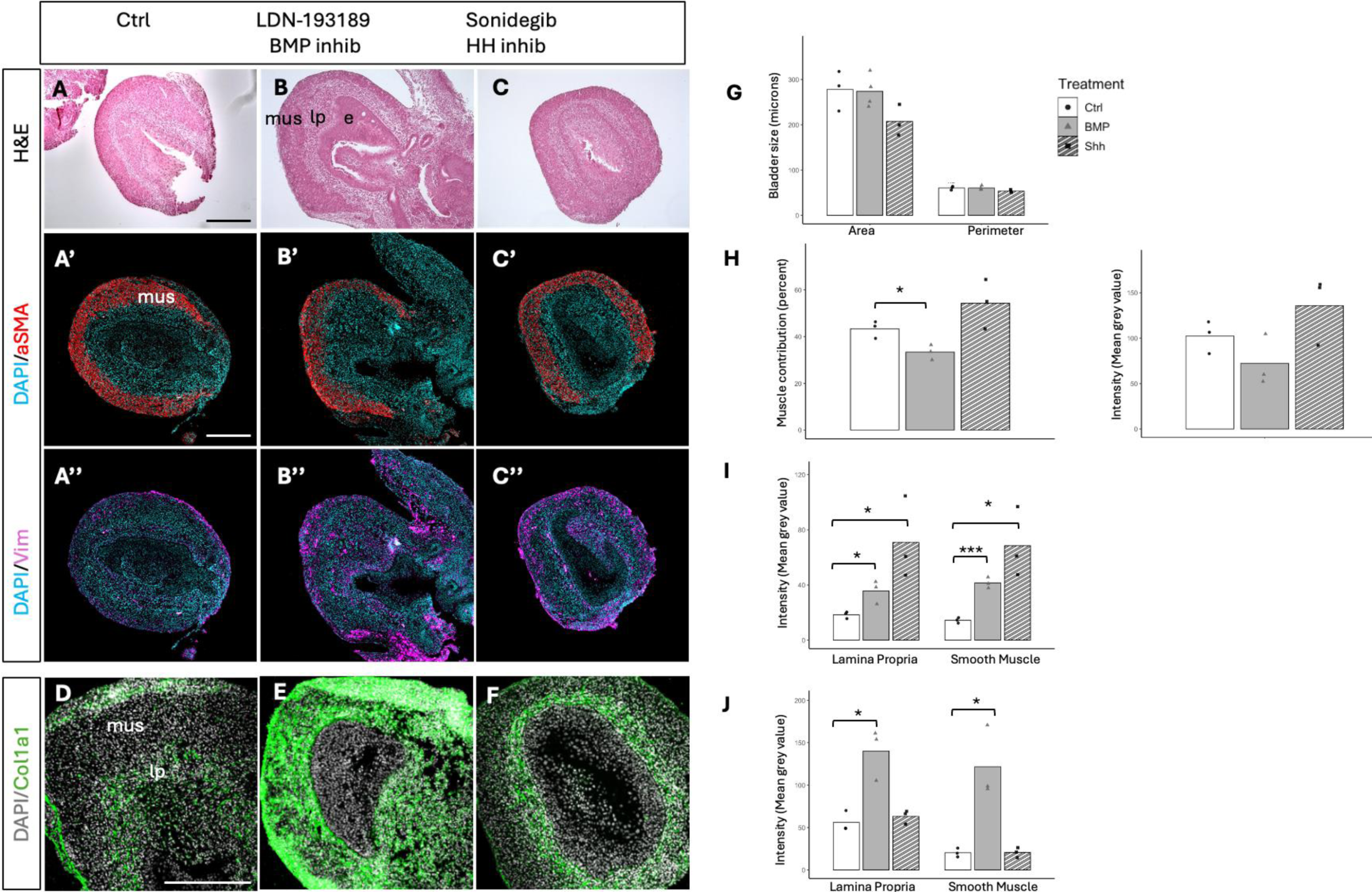
Inhibition of HH and BMP signaling disrupt bladder mesenchyme patterning. (A-J) Bladder cultures were treated with vehicle (DMSO), the BMP inhibitor LDN193189 or the HH inhibitor Sonidegib (both at 2uM). Layers indicated with e – epithelium, lp – lamina propria, mus – smooth muscle. (A-C, G) Bladder sizes were comparable between the groups with SHH-inhibited bladders showing a trend of being smaller. (A’-C’, H) Staining for aSMA revealed a significant decrease in muscle contribution to the bladder wall and a trend towards decreased alpha-smooth muscle actin expression in the muscle layer in BMP-inhibited cultures. HH-inhibited cultures showed a trend towards slightly increased muscle. (A’’ – C’’, I) Both BMP and HH-inhibited bladder cultures showed increased vimentin expression in both the lamina propria and smooth muscle layers. (D-F, J) BMP-inhibited bladders had a significant increase in col1a1 expression in the lamina propria and muscle layers. * = p<0.05, *** p < 0.001. Scale bar = 250 um.

## Discussion

To understand the molecular and cellular events during bladder development, we present the first single-cell analysis of embryonic mouse bladder development at three critical stages: E12, when the bladder first forms as a distinct outgrowth of the cloaca, E14, when the smooth muscle begins to arise, and E15, when the epithelial and mesenchymal layers of the bladder have stratified to form mature bladder tissue. Embryonic day 14 is a critical stage for cellular differentiation because many of the mesenchymal progenitor cells enter a muscle-fibroblast intermediate stage before differentiating to become smooth muscle or fibroblast cells (Figure 7). By E15, the mesenchymal progenitor population has mostly completed its differentiation into the latter two cell types. Our study demonstrates that Bmp1ra, Fgfr1, Ptch1, and Fzd2 are the key receptors that mediate BMP, FGF, HH and WNT signaling in mesenchymal progenitors, while Fgfr2 and Fzd6 mediate FGF and WNT signaling in epithelial cells. This suggests that bladder development is regulated by crosstalk between the mesenchyme and epithelium that is mediated by SHH, BMP, FGF, and WNT signaling events.

**Figure 7:**
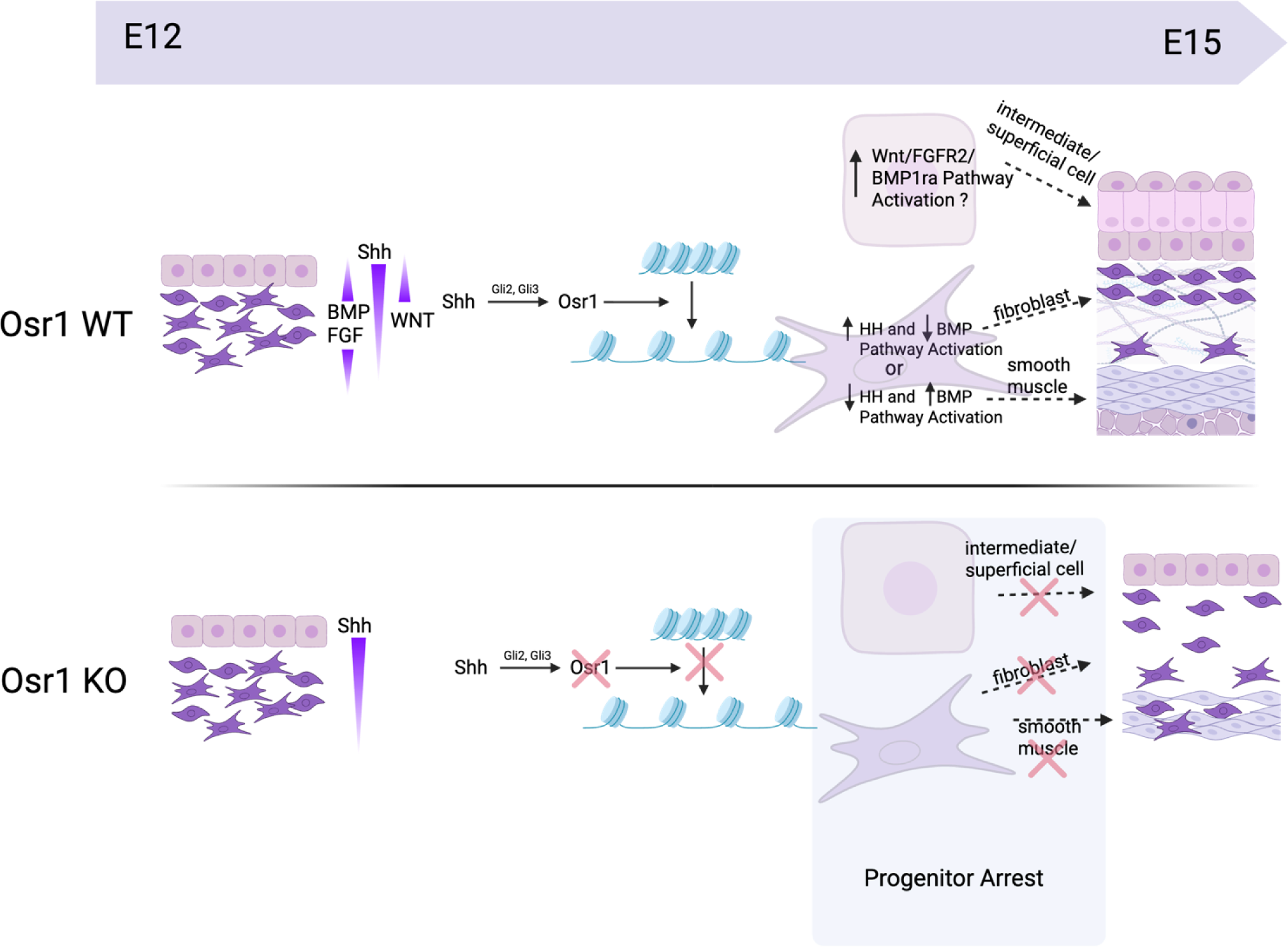
Proposed model of Osr1-mediated bladder development. Osr1 regulates effectors of bladder development within the Shh, BMP, Wnt, and FGF pathways. Receptors, ligands and transcription factors within all four pathways were downregulated at E14 when the major differentiation events occur in *Osr1*-KO mice. We propose that Osr1 acts as a pioneering transcription factor that facilitates the opening of closed chromatin regions leading to upregulation of HH, BMP, Wnt, and FGF signaling pathways and cellular differentiation. When Osr1 is not functional, progenitors remain trapped in an undifferentiated state, resulting in the loss of differentiated cell types including smooth muscle, fibroblasts, and intermediate and superficial epithelial cells.

Using inhibitors for BMP and HH, we demonstrated that these pathways are essential for bladder mesenchyme development between E13-E15. In contrast to the literature, we did not see a loss of smooth muscle differentiation with inhibition of SHH as has been shown by others (M. Cao et al., 2010; Cheng Wei et al., 2008), however this could be due to the timing of our treatment. We added inhibitors at E13, when smooth muscle development had already started. SHH is essential for proliferation and growth of the mesenchyme in the bladder and ureter, and we did see a decrease in bladder size in the cultured explants. The working model of bladder smooth muscle differentiation suggests that lower levels of HH signaling coupled with lower levels of BMP4 provide a permissive environment for smooth muscle differentiation (Liaw et al., 2018). Previous studies showed that exogenous BMP delivery inhibits smooth muscle formation, while the BMP antagonist, Noggin, increased the smooth muscle compartment (Cheng Wei et al., 2008; Huycke et al., 2019). In the ureter, BMP4 and HH both function as activators of smooth muscle differentiation, while development of the lamina propria requires other effectors including WNT and FGF signaling (Deuper et al., 2022; Straube et al., 2025). Contrary to what was seen in the gut where BMP inhibition by Noggin expanded the smooth muscle layer, BMP inhibition via LDN193189 which binds to and antagonizes Bmp1ra (the receptor for BMP4) caused a decrease in smooth muscle differentiation as well as a massive increase in collagen I deposition in the bladder. Although we used small molecule inhibitors at concentrations based on the literature, it is difficult to define the magnitude of pathway inhibition in the explants. We selected relatively high concentrations (2uM) to block the pathway without inducing tissue necrosis. Our results show that both the BMP and HH pathways play crucial roles during bladder mesenchyme patterning, and that the defects in the *Osr1-*KO embryos are due to a combination of the loss of both pathways. We speculate that Osr1 promotes mesenchymal differentiation through its activation and tight modulation of HH and BMP pathways. Smooth muscle differentiation arises from a decrease in HH activity and sustained BMP activity, while collagen-secreting fibroblast differentiation arises from increased activation of HH pathways and a decrease in BMP activity. These specific differentiation events could arise if Osr1 toggles between HH and BMP effector binding to regulate their relative expression during bladder development.

Previously our lab identified bladder defects from heterozygous loss of *Osr1*, however this model did not permit an analysis of whether Osr1 is required for bladder development. Here we show that Osr1 is essential for patterning the bladder epithelium and mesenchyme. Most strikingly, loss of *Osr1* results in the failure of the mesenchyme to differentiate into smooth muscle as well as functional fibroblasts based on the loss of collagen expression. Re-analysis of pre-existing ChIP-seq datasets shows that Osr1 binds to promoters that regulate the HH, BMP, WNT, and FGF signaling pathways. Key Osr1 binding targets include *Wnt4*, *Wnt5a*, *Fzd2*, *Ptch1*, *Smo*, *Fgf9*, *Fgfr2*, *Fgfr1*, *Smad4*, and *Smad2* that are all expressed during bladder development. A previous study also reported that Osr1 bound to and regulated *Bmp4* (Rankin et al., 2012). Furthermore, we showed that antagonism of Smo or Alk2/3 during bladder development were both able to replicate the smaller bladder and loss of smooth muscle phenotypes, respectively, observed from loss of *Osr1*. Loss of *Osr1* affected all of these signaling pathways leading to mesenchymal and epithelial progenitors that are unable to differentiate suggesting a broad role of Osr1 during bladder development (Figure 7). This is consistent with the identified role of Osr1 as a pioneer transcription factor that regulates chromatin accessibility by targeting the core histone variant protein, H2A.Z to regions of closed chromatin(Zhang et al., 2017). Although we did not evaluate H2A.Z localization in our study, we did see a decrease in H2A.Z transcripts in the *Osr1*-KO mice suggesting chromatin accessibility may be affected. Consistent with this, loss of *Osr1* results in arrest of cells in progenitor-like states in both the mesenchyme and epithelium, potentially due to the inability of Osr1 to access genes required for differentiation of the bladder.

The identified role of Osr1 in epithelial tissue is novel, as all previous studies have primarily shown its role in mesenchymal cells and their derivatives. We observed a loss of intermediate and superficial cells, while there were similar levels of basal cells. These phenotypes resemble those from previous studies where loss of *Bmp4* and *Pparg* impaired basal cell differentiation, resulting in a deficiency of superficial cells (C. Liu et al., 2019; Mysorekar et al., 2009). Superficial cells are essential for maintaining the integrity of the bladder barrier and for signaling bladder contraction, thus the *Osr1*-KO epithelial phenotype warrants further investigation. Our single-cell dataset was limited due to the relatively small numbers of epithelial cells in comparison to mesenchymal cells. In future studies, we would isolate the epithelium prior to library prep to expand these cell numbers and do a more in-depth analysis. Analysis of Osr1 binding targets by performing ChIP-seq in the embryonic bladder or a single cell ATAC-seq would further elucidate how Osr1 regulates these key signaling pathways. Another outstanding question is whether the effects on the bladder mesenchyme or epithelium from loss of *Osr1* are primary or secondary to defects in the adjacent layer, since epithelial-mesenchymal crosstalk is essential for development. Indeed, loss of signaling from the depleted suburothelial cells may result in defects in bladder epithelial stratification even without the loss of *Osr1* in the epithelia. This could be addressed by creating mesenchyme and epithelial-specific *Osr1* conditional KO mice. In conclusion, we provide a detailed outline of the key molecular events underlying early bladder development and demonstrate an essential role for Osr1 during this period. This information is an important first step in advancing bladder organoids as well as bladder tissue replacement strategies.

## Methods

### Mouse lines

*Osr1^tm1Jian^* or *Odd1-LacZ* were obtained from Jackson Laboratories and maintained on a C57BL/6J background (Jackson Lab #009387). These mice contain a β-galactosidase reporter fused to an *Osr1* null allele. These mice were compared to Osr1^+/+^ littermates in all analyses. Hoxb7-GFP CD1 mice were used for the bladder culture experiments. Timed matings were set up in the late afternoon with plug checks every morning with T0 assumed to be midnight on the day of a positive plug. All mice were housed in ventilated cages with up to five mice/cage with wood chip bedding and a diurnal light cycle providing 12 h of light. The mice fed and drank from their water bottles ad libitum with feed provided as per the US National Research Council recommendations for rodent nutrition. Environmental enrichment with either paper strands or cellulose-based shelters was included in each cage. The room temperature was maintained between 18 and 24°C, and the humidity was maintained between 30 and 70%. All animal studies were performed in accordance with the regulations of the Canadian Council on Animal Care and approved by the McGill University Animal Care Committee (AUP 4120).

### Tissue Collection and paraffin Embedding

Mice were euthanized at E12, E14, and E15 and staged using crown-rump measurements. Lower torsos of embryos were dissected into cold PBS and fixed in 4% PFA overnight for sectioning, while limbs were taken for DNA extraction and genotyping of the *Osr1-LacZ* allele. Fixed tissues were then transferred to 70% ethanol for long term storage at 4°C. Some tissues were processed for paraffin embedding as previously described. Bladders were sectioned at 5um thickness.

### RNAScope for mRNA expression

Using the multiplex fluorescent V2 kit and the 4 plex ancillary kit [RNAScope, ACD Biosystems, CA] mRNA detection was performed for *acta2* for alpha-smooth muscle actin, and *osr1* for Odd-skipped related 1. Staining was performed according to the manufacturer’s directions.

### Immunofluorescent staining

Paraffin sections were deparaffinized and rehydrated in PBS. Heat-mediated citrate antigen retrieval was performed, and sections were blocked in 1% BSA for 1 hour before proceeding to staining. Primary antibody incubation was performed at 4°C overnight for rabbit anti-mouse alpha-smooth muscle actin (Abcam) at a dilution of 1/100, mouse anti-mouse vimentin (Santa Cruz) at a dilution of 1/100, goat anti-mouse Col1a1 (Southern Biotech) at a dilution of 1/250, and mouse AE1/3 monoclonal anti-keratin antibodies kind gift from Dr. TT Sun (Cooper & Sun, 1986) at a dilution of 1/5. Sections were then washed and incubated with secondary antibodies (Invitrogen) as follows, anti-rabbit 488, anti-mouse 555, or anti-goat 488 for 1 hour at room temperature at 1/500 dilution. Sections were then washed and mounted using the antifade reagent (prolong gold). Sections were imaged under confocal microscopy (Leica LSM880 or LSM780). Image processing was done using the Zen black software and Fiji. Intensity was measured using mean grey value of evenly sized and spaced sampling squares throughout the bladder wall in the lamina propria and muscle separately and significance was evaluated using a two-tailed students t-test.

### Single-cell sample preparation

Bladders were dissected from E12, E14 and E15 embryos (staged by crown-rump measurements) on ice, as depicted in figure 1A, and limbs were taken from each embryo for DNA extraction and genotyping. Bladders were then stored in Hypothermosol (Stem cell Technologies) on ice to preserve cells and mRNA while genotyping was performed. 2 bladders from each genotype were pooled for each timepoint to average out biological variability. Tissue was digested by immersion in a solution containing 1mg of protease from *B.licheniformis* (Sigma) in 10ul of DPBS, 45ul of Accutase and 45 ul of Accumax (Stem cell Technologies) solution as previously described (Sekiguchi & Hauser, 2019)). Dissociation was halted using 10% FBS in DPBS and samples were passed through Flowmi cell strainers (40um) into fresh Eppendorf tubes before being sent to the McGill Genome Center for quality control and cell viability assessment and library preparation using the 10x Genomics NextGem scRNA 3’ V3.1. The libraries were then converted into MGI and sequenced using MGI DNBSEQ G-400, PE100.

### Single-cell sequencing analysis

Sequencing alignments and sample assessments were done using cellRanger performed by the McGill Genome Center before filtered feature and raw feature matrix H5 files were transferred to us for analysis. Data was in R version 4.4.0 and converted into Seurat V5 objects. Standard pre-processing and quality control was performed using the ‘subset’ command to only keep cells with feature counts between 200 and 3000 and mitochrondrial percentage below 15. Data was normalized using SCTransform, and LogNormalize at a scale factor of 10,000. Data across timepoints and genotypes was integrated using Harmony v0.1 after integration features were selected (3000) and PCA dimensionality reduction (50 principal components) was performed. UMAPs were generated using runUMAP (40 dimensions) and clusters were identified using FindNeighbours and FindClusters. Resolution was adjusted between 0.1 and 0.4 depending on silhouette scores. Clusters were characterized using canonical cell type markers. Cell proportions within clusters were compared across timepoints and genotypes. Pseudotime Lineage prediction was performed using Monocle 3.0. Pseudobulk differential gene expression analysis was performed using FindMarkers with log fold change threshold set to 0.25 and assessed for significance using the default Wilcoxon rank sum test. Significance was set as FDR p value < 0.05. Gene ontology analysis was performed using ClusterProfiler and the org.Mm.eg.db database and pathways corresponding to upregulated and downregulated genes in the knockout samples were generated.

### Bladder culture assays

Bladders were dissected from E13 mouse embryos and stored in cold PBS on ice. Bladders were then cultured using an air-liquid interface culture system. Briefly, cell culture inserts (0.4 um Millicell PTFE standing cell culture inserts (Sigma)) were placed into 6 well plates containing 1.2ml of warmed DMEM/F12 media supplemented with insulin-transferrin-selenium solution (1x) and pen-strep (1x) with vehicle (DMSO), 2 uM SHH inhibitor (Sonidegib) or 2 uM BMP inhibitor (LDN193189) from Medchemexpress. Bladders were then placed on cell culture inserts and cultured in incubators set to 37 °C and 5% CO2. Bright field imaging was performed daily to ensure continued growth and bladders were removed from the insert membrane and placed into 4% PFA and processed for histology as previously mentioned.

## Supplemental Material

**Supplemental Table 1:** Single cell sequencing sample preparation summary.

| Sample | Number<br>pooled | Cells/ul | Total cells | Viability<br>(%) | Cells used for<br>capture |
| --- | --- | --- | --- | --- | --- |
| E12 WT | 2 | 1200 | 36,000 | 98 | 16,000 |
| E12 KO | 2 | 400 | 18,000 | 87 | 13,000 |
| E14 WT | 2 | 2600 | 78,000 | 97 | 16,000 |
| E14 KO | 2 | 2100 | 84,000 | 96 | 16,000 |
| E15 WT | 1 | 800 | 40,000 | 98 | 16,000 |
| E15 KO | 1 | 500 | 25,000 | 99 | 16,000 |

**Supplemental Figure 1:**
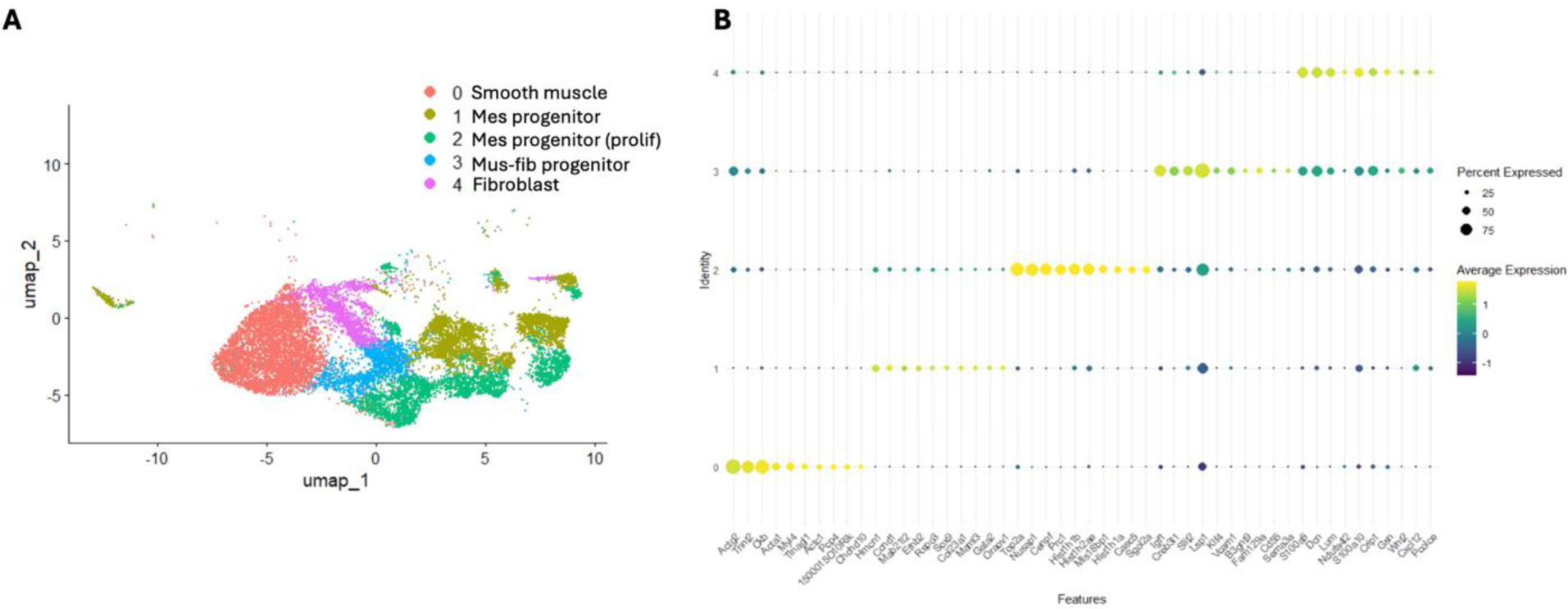
Markers for mesenchymal cell populations in WT embryonic bladders integrated from E12 to E15. (A) UMAP of cell populations, Mes – mesenchymal, mus – muscle, fib – fibroblast (B) Top 10 markers obtained through FindAllMarkers function in Seurat.

**Supplemental Figure 2:**
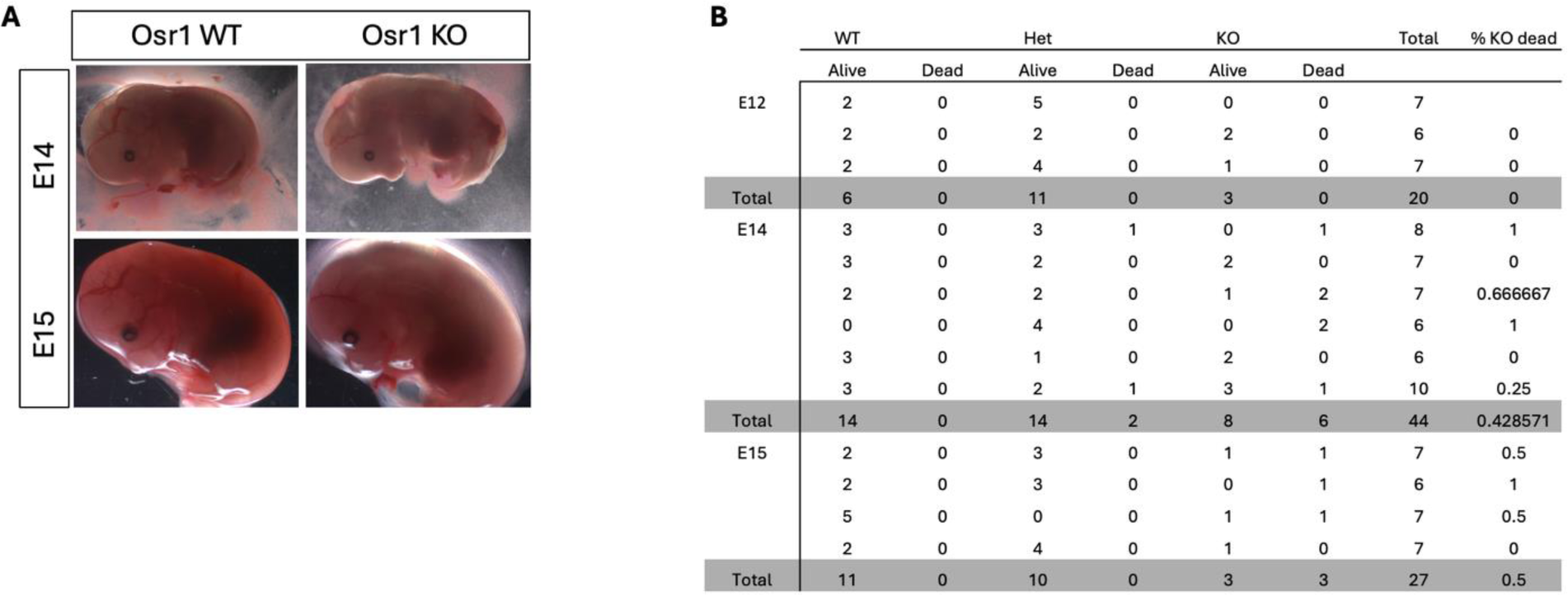
Summary of breeding and embryo viability from Osr1 het by het cross: (A) Representative whole mount imaging of E14 and E15 embryos with noticeable spinal edema seen in KO embryos. (B) Summary of viable vs dead embryos, with approximately 50% of *Osr1*-KO embryos surviving until E15. KO mice were not viable at later timepoints.

**Supplemental Figure 3:**
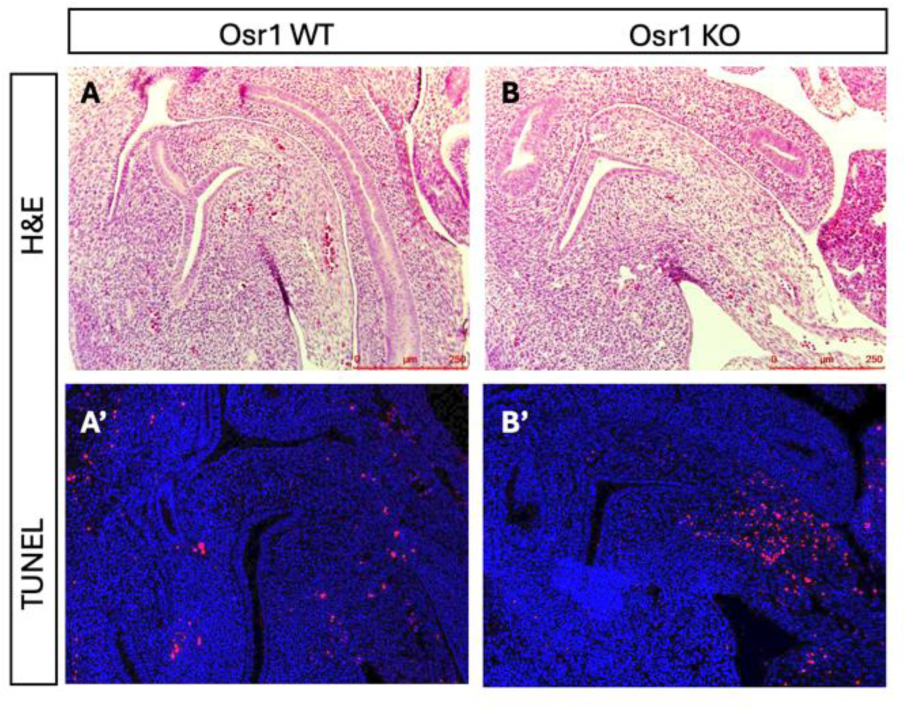
TUNEL cell death analysis at E12: (A) WT embryos had very little cell death with some occurring at the edge of the putative bladder (B) KO embryos displayed higher levels of cell death in a similar region to the WT embryos but at higher levels. In both the WT and KO embryos, cell death was concentrated in a region close to the umbilical artery and away from the cloacal epithelium suggesting this region may not represent the future bladder.

**Supplemental Figure 4:**
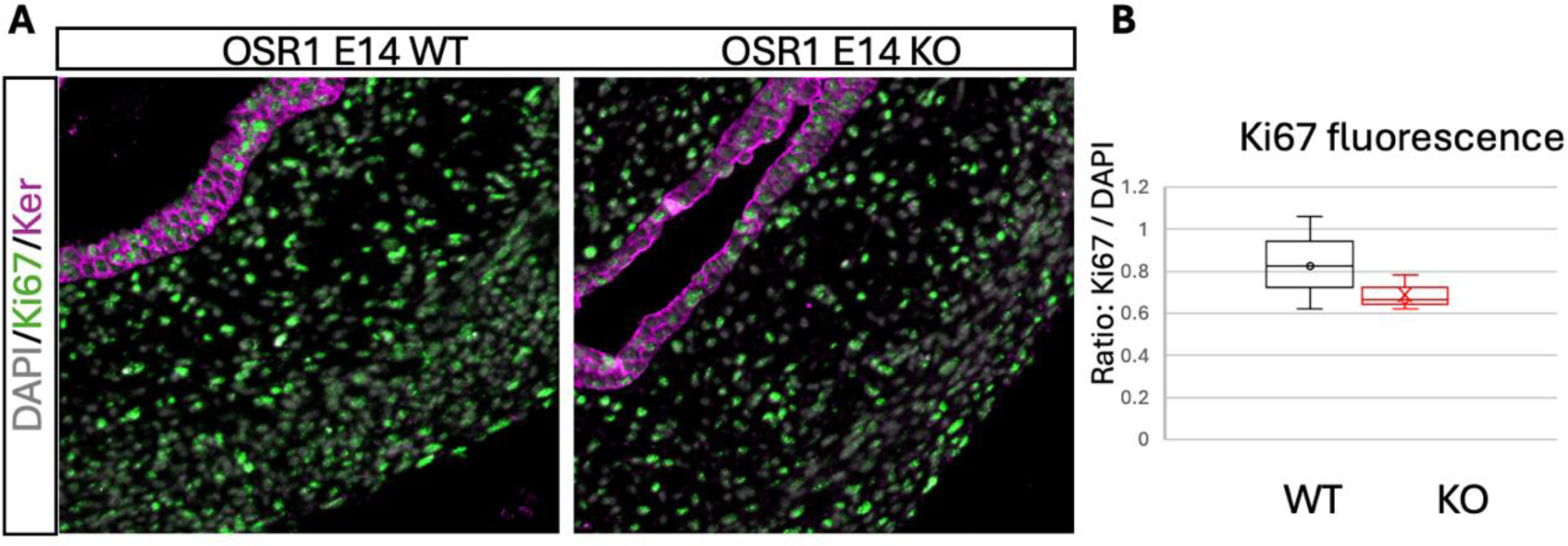
Ki67 proliferation analysis at E14: (A) Both WT and KO embryos had high levels of cell proliferation with most mesenchymal and epithelial cells (labelled in magenta for keratin) positive for Ki67 (green). (B) Quantification of Ki67 fluorescence normalized to DAPI fluorescence, n = 3 bladders, p = 0.3

**Supplemental Figure 5:**
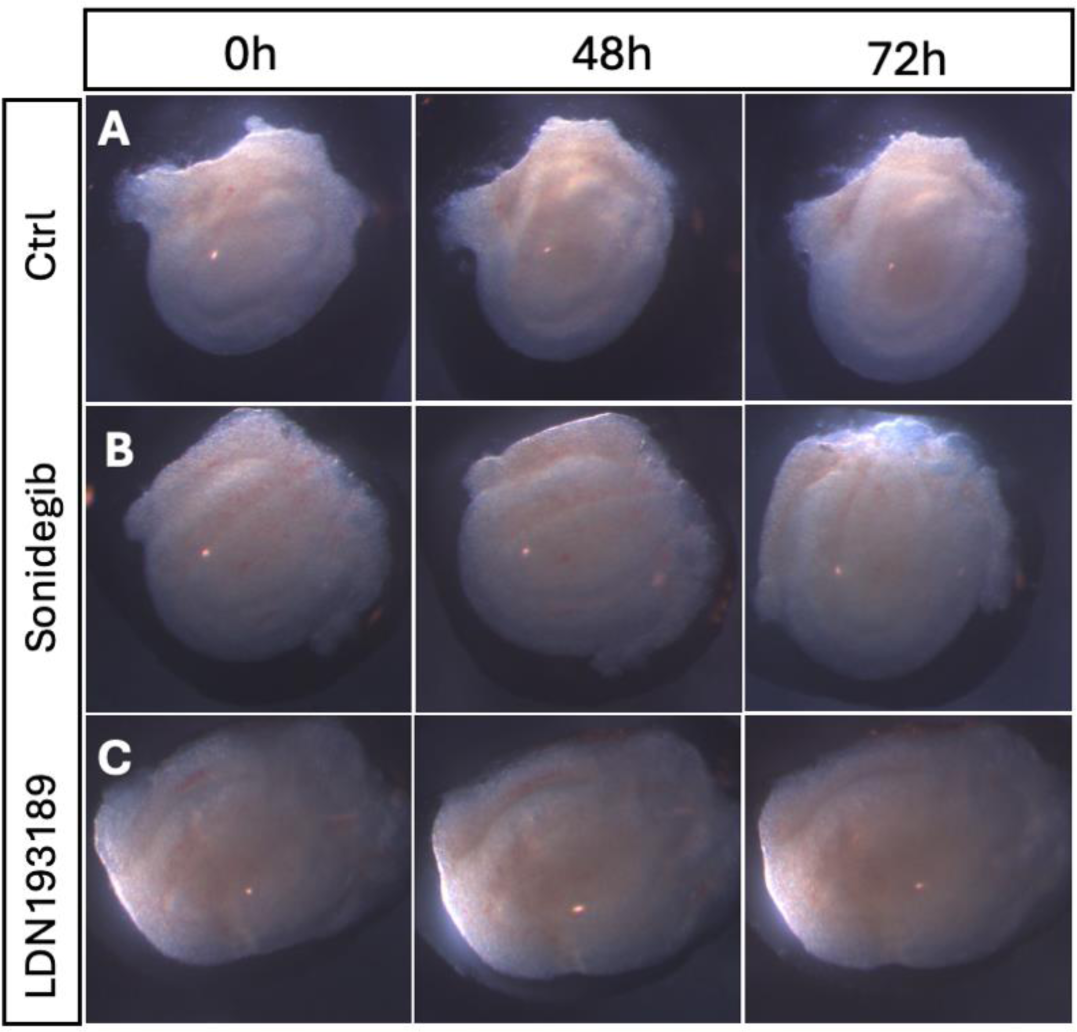
Representative whole mount imaging of bladder cultures taken at 0, 48 and 72 hours during treatment with inhibitors (A) Treated with DMSO control (B) Treated with the Smo inhibitor Sonidegib to block the SHH pathway and (C) Treated with the Alk2/3 inhibitor LDN193189 to block the BMP signaling pathway.

