## supplemental figures for "The transcription factor Osr1 regulates epithelial-mesenchymal crosstalk during embryonic bladder development"

### Slide 1
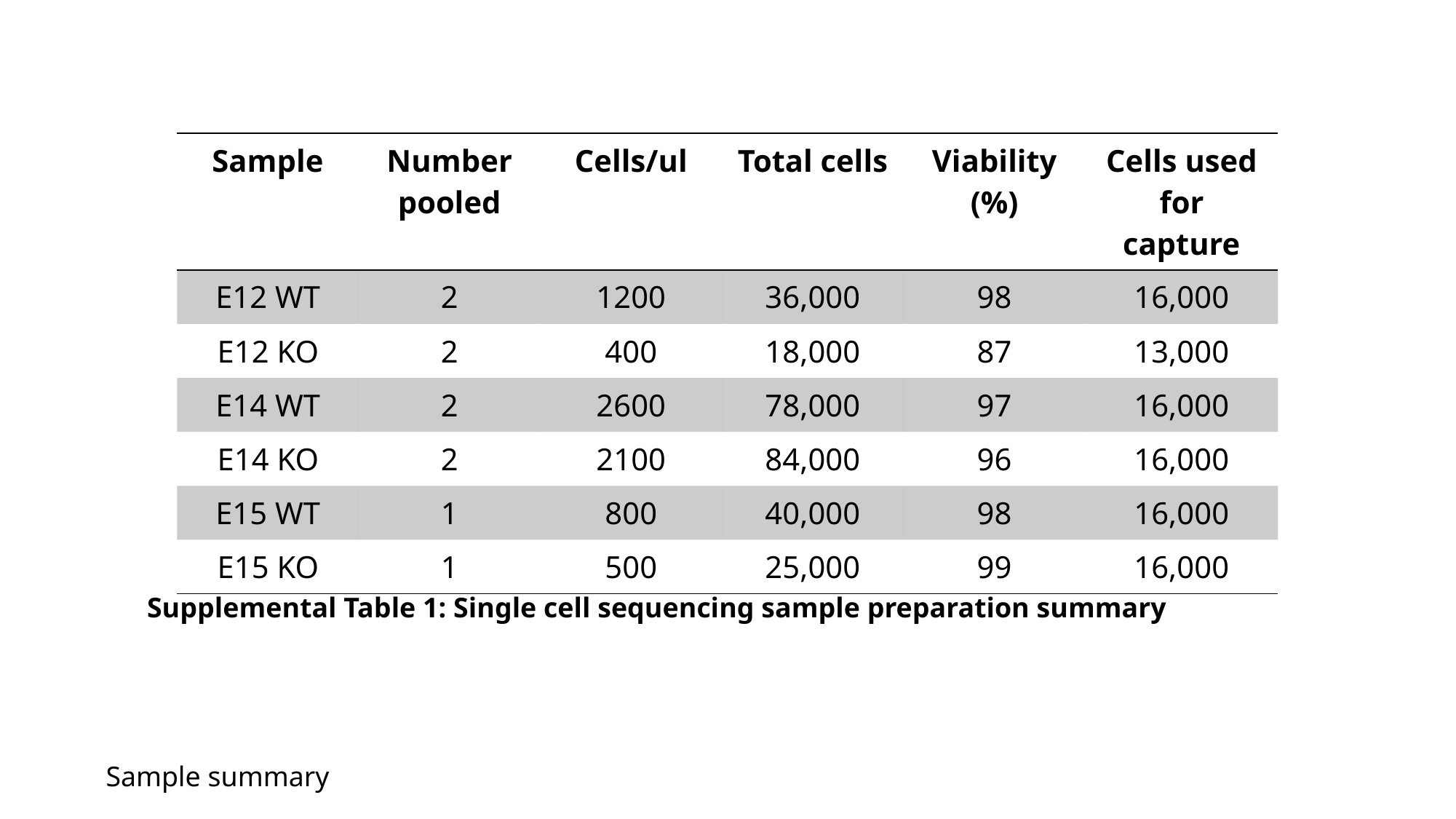

| Sample | Number pooled | Cells/ul | Total cells | Viability (%) | Cells used for capture |
| --- | --- | --- | --- | --- | --- |
| E12 WT | 2 | 1200 | 36,000 | 98 | 16,000 |
| E12 KO | 2 | 400 | 18,000 | 87 | 13,000 |
| E14 WT | 2 | 2600 | 78,000 | 97 | 16,000 |
| E14 KO | 2 | 2100 | 84,000 | 96 | 16,000 |
| E15 WT | 1 | 800 | 40,000 | 98 | 16,000 |
| E15 KO | 1 | 500 | 25,000 | 99 | 16,000 |
Supplemental Table 1: Single cell sequencing sample preparation summary
Sample summary

### Slide 2
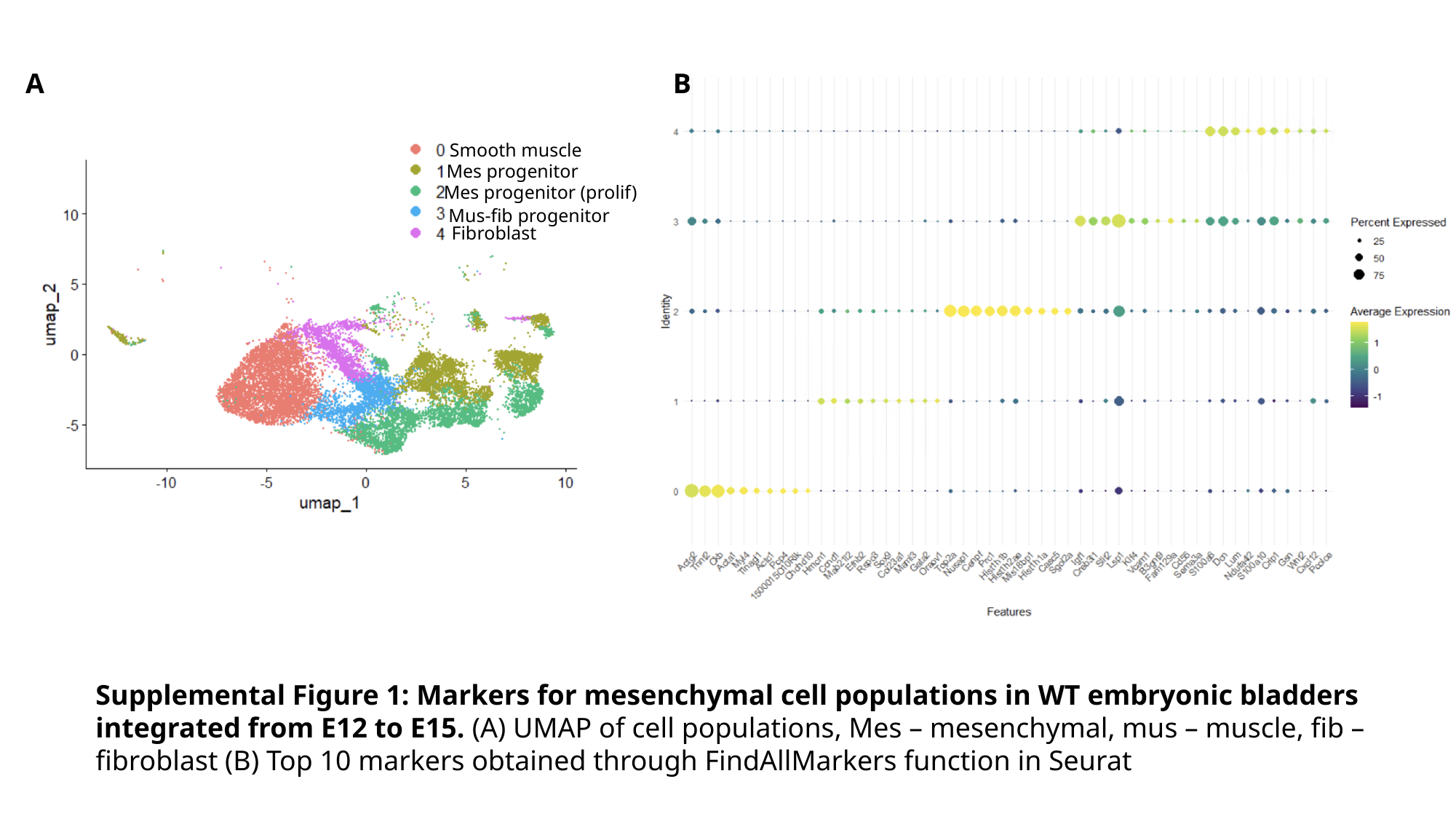

A
B
Smooth muscle
Mes progenitor
Mes progenitor (prolif)
Mus-fib progenitor
Fibroblast
Supplemental Figure 1: Markers for mesenchymal cell populations in WT embryonic bladders integrated from E12 to E15. (A) UMAP of cell populations, Mes – mesenchymal, mus – muscle, fib – fibroblast (B) Top 10 markers obtained through FindAllMarkers function in Seurat

### Slide 3
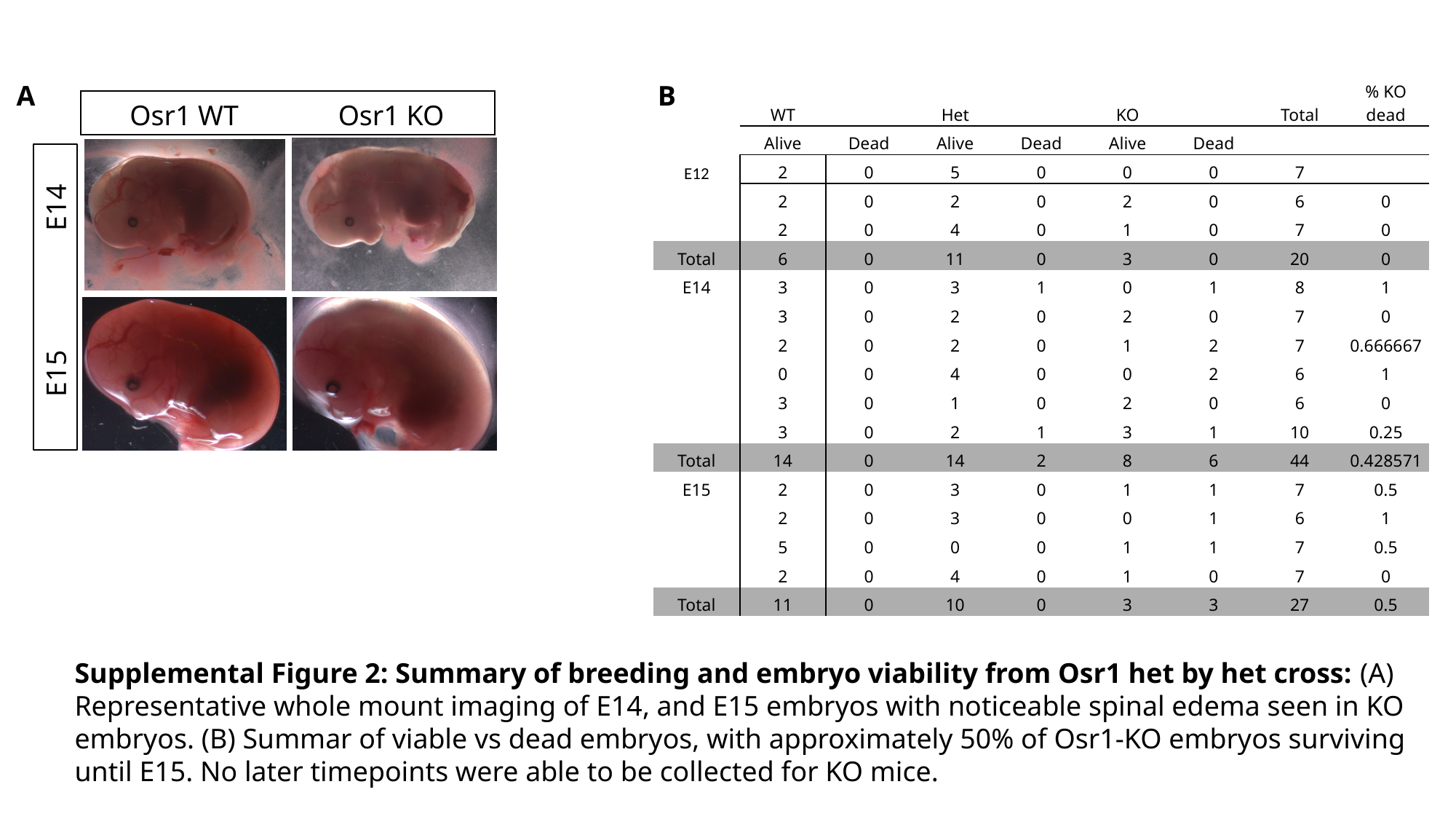

A
B
| | WT | | Het | | KO | | Total | % KO dead |
| --- | --- | --- | --- | --- | --- | --- | --- | --- |
| | Alive | Dead | Alive | Dead | Alive | Dead | | |
| E12 | 2 | 0 | 5 | 0 | 0 | 0 | 7 | |
| | 2 | 0 | 2 | 0 | 2 | 0 | 6 | 0 |
| | 2 | 0 | 4 | 0 | 1 | 0 | 7 | 0 |
| Total | 6 | 0 | 11 | 0 | 3 | 0 | 20 | 0 |
| E14 | 3 | 0 | 3 | 1 | 0 | 1 | 8 | 1 |
| | 3 | 0 | 2 | 0 | 2 | 0 | 7 | 0 |
| | 2 | 0 | 2 | 0 | 1 | 2 | 7 | 0.666667 |
| | 0 | 0 | 4 | 0 | 0 | 2 | 6 | 1 |
| | 3 | 0 | 1 | 0 | 2 | 0 | 6 | 0 |
| | 3 | 0 | 2 | 1 | 3 | 1 | 10 | 0.25 |
| Total | 14 | 0 | 14 | 2 | 8 | 6 | 44 | 0.428571 |
| E15 | 2 | 0 | 3 | 0 | 1 | 1 | 7 | 0.5 |
| | 2 | 0 | 3 | 0 | 0 | 1 | 6 | 1 |
| | 5 | 0 | 0 | 0 | 1 | 1 | 7 | 0.5 |
| | 2 | 0 | 4 | 0 | 1 | 0 | 7 | 0 |
| Total | 11 | 0 | 10 | 0 | 3 | 3 | 27 | 0.5 |
| Osr1 WT | Osr1 KO |
| --- | --- |
 E15	 E14
Supplemental Figure 2: Summary of breeding and embryo viability from Osr1 het by het cross: (A) Representative whole mount imaging of E14, and E15 embryos with noticeable spinal edema seen in KO embryos. (B) Summar of viable vs dead embryos, with approximately 50% of Osr1-KO embryos surviving until E15. No later timepoints were able to be collected for KO mice.

### Slide 4
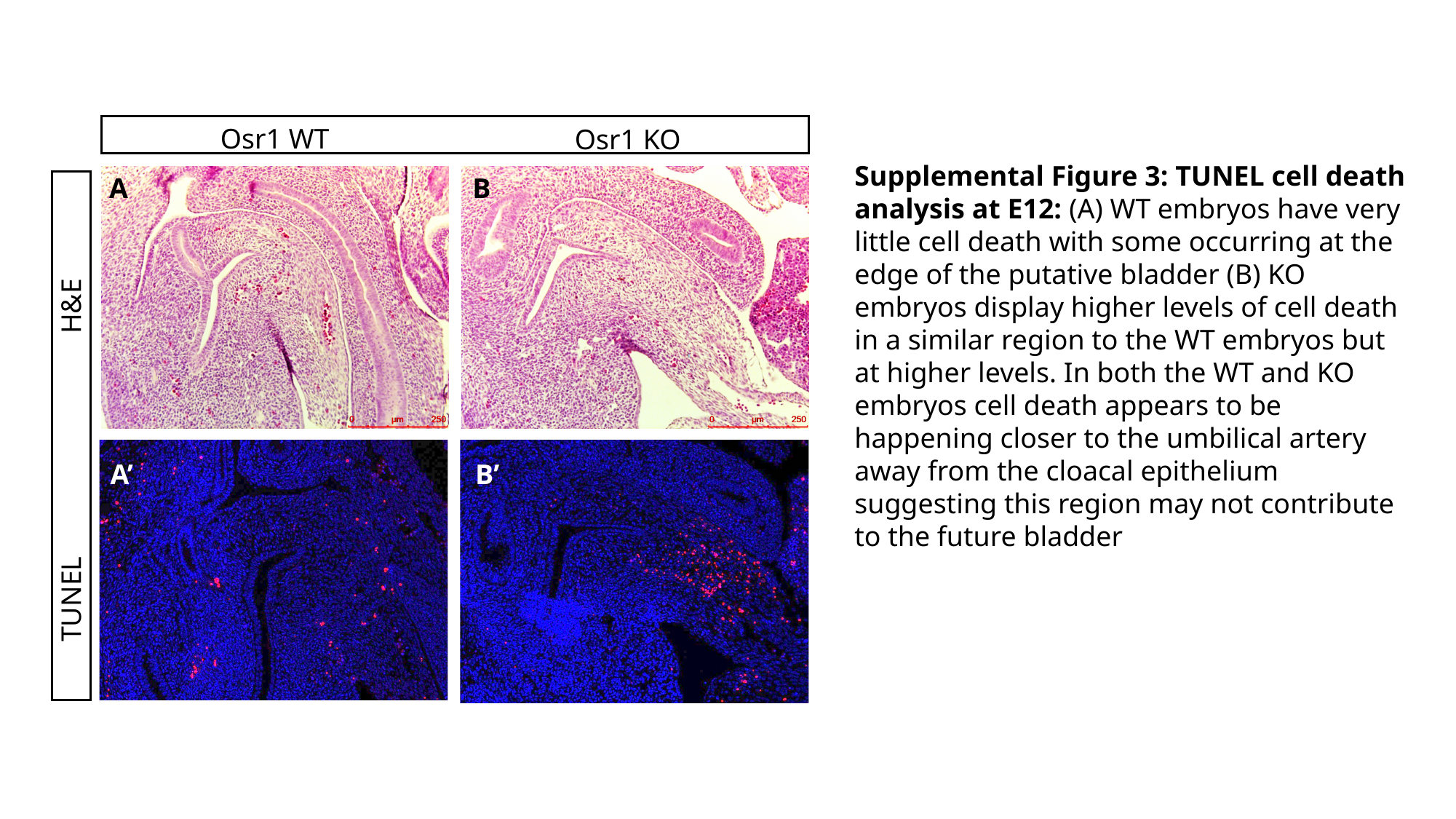

Osr1 WT
Osr1 KO
A
B
H&E
A’
B’
TUNEL
Supplemental Figure 3: TUNEL cell death analysis at E12: (A) WT embryos have very little cell death with some occurring at the edge of the putative bladder (B) KO embryos display higher levels of cell death in a similar region to the WT embryos but at higher levels. In both the WT and KO embryos cell death appears to be happening closer to the umbilical artery away from the cloacal epithelium suggesting this region may not contribute to the future bladder

### Slide 5
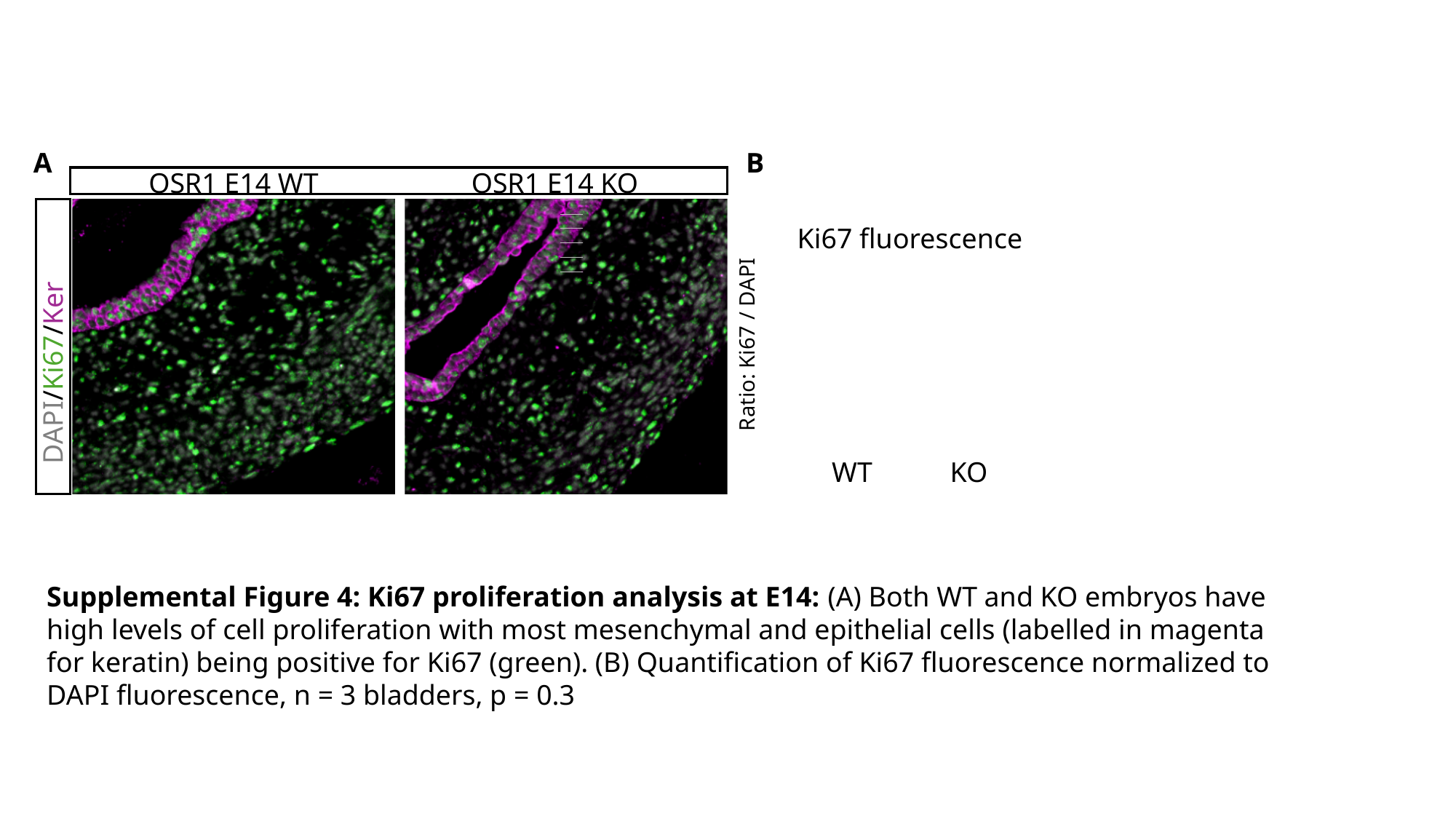

A
B
OSR1 E14 WT
OSR1 E14 KO
Ki67 fluorescence
Ratio: Ki67 / DAPI
DAPI/Ki67/Ker
WT KO
Supplemental Figure 4: Ki67 proliferation analysis at E14: (A) Both WT and KO embryos have high levels of cell proliferation with most mesenchymal and epithelial cells (labelled in magenta for keratin) being positive for Ki67 (green). (B) Quantification of Ki67 fluorescence normalized to DAPI fluorescence, n = 3 bladders, p = 0.3

### Slide 6
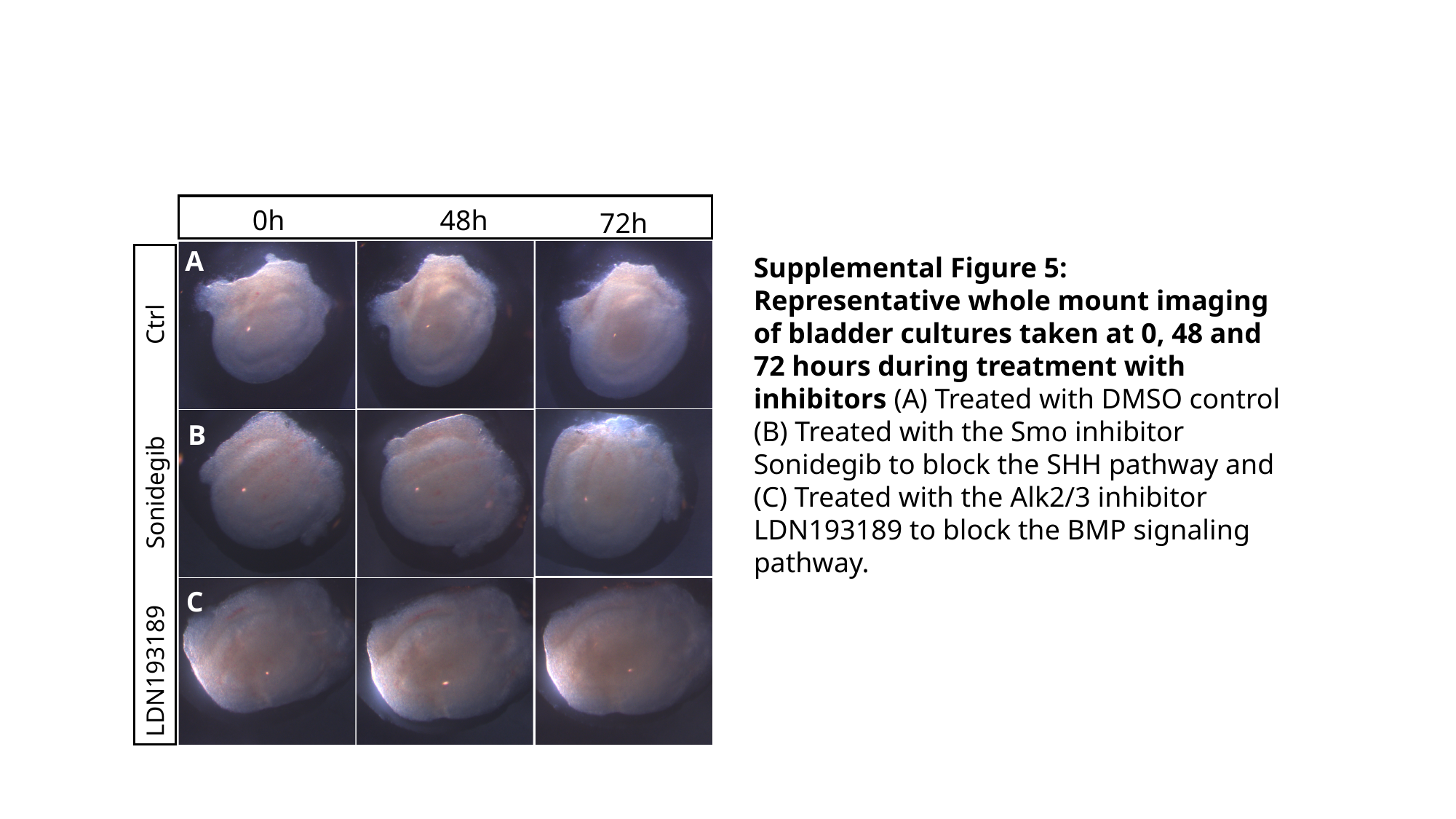

0h
48h
72h
A
Supplemental Figure 5: Representative whole mount imaging of bladder cultures taken at 0, 48 and 72 hours during treatment with inhibitors (A) Treated with DMSO control (B) Treated with the Smo inhibitor Sonidegib to block the SHH pathway and (C) Treated with the Alk2/3 inhibitor LDN193189 to block the BMP signaling pathway.
Ctrl
B
Sonidegib
C
LDN193189
